# Cortical source imaging of resting-state MEG with a high resolution atlas: An evaluation of methods

**DOI:** 10.1101/2020.01.12.903302

**Authors:** Luke Tait, Ayşegül Özkan, Maciej J Szul, Jiaxiang Zhang

## Abstract

Non-invasive functional neuroimaging of the human brain at rest can give crucial insight into the mechanisms that underpin healthy cognition and neurological disorders. Magnetoencephalography (MEG) measures extracranial magnetic fields originating from neuronal activity with very high temporal resolution, but requires source reconstruction to make neuroanatomical inferences from these signals. Many source reconstruction algorithms for task-based MEG data are available. However, no consensus yet exists on the optimum algorithm for resting-state data.

Here, we evaluated the performance of six commonly-used source reconstruction algorithms based on minimum-norm and beamforming estimates. In the context of human resting-state MEG, we compared the algorithms using quantitative metrics, including resolution properties of inverse solutions and explained variance in sensor-level data. Next, we proposed a data-driven approach to reduce the atlas from the Human Connectome Project’s multimodal parcellation of the human cortex. This procedure produced a reduced cortical atlas with 250 regions, optimized to match the spatial resolution and the rank of MEG data from the current generation of MEG scanners.

For both voxel-wise and parcellated source reconstructions, we showed that the eLORETA inverse algorithm had zero localization error, high spatial resolution, and superior performance in predicting sensor-level activity. Our comprehensive comparisons and recommandations can serve as a guide for choosing appropriate methodologies in future studies of resting-state MEG.

## 1. Introduction

Non-invasive imaging of the human brain’s restingstate dynamics can give insight into the neuronal mechanisms underpinning healthy cognition and neurological disorders (van den Heuvel and Hulshoff Pol, 2010; Douw et al., 2011; Babiloni et al., 2016; Cohen, 2018; Michel and Koenig, 2018; Lee et al., 2019). Haemodynamic or metabolic correlates of resting-state neuronal activity such as functional MRI (fMRI) (Greicius et al., 2003) or positron emission tomography (Raichle et al., 2001) have identified key brain networks exhibiting spontaneous functional connectivity during rest, but their temporal resolutions are much slower than the timescale of neuronal oscillations. Hence, properties such as oscillatory power spectra and phase-binding (Babiloni et al., 2016), and the brain’s rapid transitions in active networks at the temporal order of tens to hundreds of milliseconds (Michel and Koenig, 2018) cannot be studied using these modalities. Conversely, conventional magneto-or electro-encephalography (M/EEG) measure macroscopic neuronal post-synaptic activity at a high temporal resolution of the order of milliseconds, but sensors are located outside of the scalp and cannot directly infer the spatial origin of signals.

To make neuroanatomical inferences on the M/EEG signal, source reconstruction must be used (Michel et al., 2004; Michel and Brunet, 2019). This problem is ill-posed and therefore has no unique solution, but the neurophysiologically-informed assumptions underpinning M/EEG signals can be used to constrain the solution. As a result, a wide range of algorithms exist to solve this inverse problem based on different assumptions and priors (Hämäläinen and Ilmoniemi, 1994; Pascual Marqui et al., 1994; Van Veen et al., 1997; Huang et al., 1998; Mosher and Leahy, 1998; Fuchs et al., 1999; Dale et al., 2000; Pascual-Marqui, 2002, 2007; Friston et al., 2008). Much effort has been made to assess the quality of source reconstruction algorithms in the literature, which mainly focuses on source *localization*, i.e. identifying the origin of a small number of sources, for example evoked potentials with ground truth based on known task-relevant activity (Bai et al., 2007; Hassan et al., 2014; Seeland et al., 2018; Halder et al., 2019), or simulated activity at a small number of dipoles (Bradley et al., 2016; Finger et al., 2016; Barzegaran and Knyazeva, 2017; Hassan et al., 2017; Hincapié et al., 2017; Bonaiuto et al., 2018; Pascual-Marqui et al., 2018; Anzolin et al., 2019; Halder et al., 2019).

Two issues remain unsolved in applying source reconstruction algorithms to resting-state MEG data. First, resting-state data differs from task evoked data in that activity is widely distributed across the cortex (Smith et al., 2009) and differs in signal-to-noise ratio (SNR) due to the inability to average over multiple trials (Parkkonen, 2010). Hence an algorithm optimized for task-based source localization is not necessarily optimal for resting-state source reconstruction, and it is currently no consensus on which algorithm to use for source space reconstruction of resting-state dynamics. Here, we address this question by analysing the variance of sensor space data explained by the source estimates and by analysing theoretical resolution properties (Hauk et al., 2019) of six widely used algorithms: the linearly constrained minimum variance (LCMV) beamformer (Van Veen et al., 1997) and its depth normalized counterpart (weighted LCMV, Hillebrand et al., 2012), three methods based on least-squares minimum norms under different prior assumptions of source covariance, namely the minimum norm estimate (MNE) (Hämäläinen and Ilmoniemi, 1994), weighted MNE (Fuchs et al., 1999; Lin et al., 2006), and eLORETA (Pascual-Marqui, 2007, 2009)), as well as a variance-normalized MNE, sLORETA (Pascual-Marqui, 2002).

Second, source localization methods are commonly conducted at the vertex level. It is typical in the analysis of spatially distributed resting-state data to parcellate the source-localized data into a small number of regions of interest (ROIs) for measuring functional connectivity (Tewarie et al., 2014; Colclough et al., 2016; Brookes et al., 2016; Dauwan et al., 2019). One commonly used template for cortical parcellation is the AAL atlas (Tzourio-Mazoyer et al., 2002), which is based on anatomical landmarks. Recently, there is a surge of interest in applying the Human Connectome Project Multimodal Parcelation (HCP-MMP) atlas (Glasser et al., 2016) for parcellation of cortical dynamics (Ito et al., 2017; Dubois et al., 2018; Dermitaş et al., 2019; Preti and Van De Ville, 2019; Watanabe et al., 2019). The ROI boundaries in the HCP-MMP atlas are defined by combined characteristics in cortical architecture, function, connectivity, and topography, offering better neuroanatomical precision and functional segregation. However, the HCP-MMP atlas consists of 360 cortical ROIs, and the use of such a high resolution atlas is problematic for MEG due to its limited spatial resolution. For example, the current generation of MEG scanners has 200-300 sensors (McCubbin et al., 2004). Hence, parcellation of source space data into more than this number of ROIs results in rank deficiency. This is particularly an issue if one aims to measure functional networks, since multivariate orthogonalization, which is often used to reduce spurious correlations due to source leakage (Colclough et al., 2015), requires full rank data. Here, we present a reduction of the HCP-MMP atlas with 250 ROIs, using a data-driven method based upon the forward transformation between source dynamics and the MEG sensors. We extend further our variance explained and resolution analyses to parcellated data, to assess source localization algorithms appropriate for ROI-based resting-state analyses.

The results of this study could serve as a guide for (1) choosing appropriate source reconstruction algorithms for analysis of resting-state MEG, and (2) deriving appropriate atlases for parcellation of resting-state MEG functional dynamics and connectivity. To facilitate these contributions, we have made our analysis scripts and the MEG-optimized HCP-MMP atlas open-source and freely available (https://github.com/lukewtait/evaluate_inverse_methods).

## 2. Materials and methods

### 2.1. Source reconstruction algorithms

Many methods exist for source reconstructing M/EEG data. All methods discussed in the current study involve inverting a *forward model*,

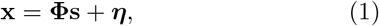

where 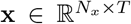 is the empirical M/EEG data (here, *N_x_* is the number of gradiometers/electrodes, and *T* is the number of sampling points), 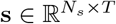 is the source data (here, *N_s_* is the number of dipoles), *η* is a noise term, and 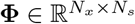 is the scalar leadfield matrix (i.e. dipole orientations are known). Details of leadfield matrix construction are outlined in section 2.6, and a more general review of methods is given by Hallez et al. (2007). For resting-state data, dipoles are generally distributed evenly across the grey matter of the cortical surface to construct the leadfield (Dale et al., 2000; Hillebrand and Barnes, 2003). Source data can then be estimated as

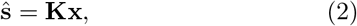

where 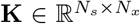 is an inverse matrix, often called the spatial filter. Algorithms to estimate **K** can be separated into beamforming and least-squares minimum-norm type estimates. Below, we briefly introduce the algorithms considered in the current study. The mathematical details of each of these solutions are provided in Appendix A.

Beamforming involves scanning over each dipole in the source space individually, calculating the corresponding row of the spatial filter independently of all other dipoles. The linearly constrained minimum-variance (LCMV) beamformer (Van Veen et al., 1997) requires the solution has unit output (i.e. all diagonals of **KΦ** are one), and that cross talk from other locations is minimized (i.e. non-diagonals of **KΦ** are minimized). This filter is known to mislocalize deep sources to superficial source points. To account for this, the beamformer weights can be normalized to unit vector norm (Hillebrand et al., 2012). Here, we refer to this solution as the *weighted LCMV* (wLCMV).

Whilst beamformers consider each source point individually, regularized minimum norm type solutions reconstruct all sources within the source space simultaneously, by minimizing the difference between the data **x** and the predicted data **Φŝ**, subject to regularization. A number of solutions have been proposed, differing in the prior estimate of source covariance, **W**. The simplest solution, often known as the minimum norm estimate (MNE; Hämäläinen and Ilmoniemi (1994)), sets **W** = **I**. Much like the LCMV beamformer, the MNE solution suffers from depth bias, so the *weighted minimum norm estimate* (wMNE) sets the diagonals of **W** inversely proportional to the sum of the leadfield (Fuchs et al., 1999; Lin et al., 2006), essentially assuming *a priori* that sources which only weakly influence the M/EEG must have a higher variance to be picked up by the sensors. The *exact low resolution electromagnetic tomography* (eLORETA) solution (Pascual-Marqui, 2007, 2009) extends this assumption further, using an algorithm to optimise the diagonals of **W** such that not only is the depth bias accounted for, the solution attains theoretically exact localization (Pascual-Marqui, 2007).

Finally, *standardized low resolution electromagnetic tomography* (sLORETA; Pascual-Marqui (2002)), an alternative approach to account for depth bias in the source localization, is to estimate *standardized* distributions of current density normalized by expected variance of each source. sLORETA first calculates the MNE solution, and then normalizes each time point to unit theoretical variance. This standardization reduces the depth bias and also has theoretically exact localization, but the resulting solution is no longer a measure of current density, and is more appropriately interpreted as the probability of source activation.

Note that all the six algorithms considered here make prior estimates of source variance, but make no prior estimates of source covariance, since the matrices **W** are always diagonal. For all source reconstruction algorithms, we used the implementations in Fieldtrip (Oostenveld et al. (2011); http://www.ru.nl/neuroimaging/fieldtrip, version 201907-16).

### 2.2. Comparing algorithms

We used two key analyses to compare source reconstruction algorithms (Figure 1). In the variance explained analysis, we estimate the variance of the empirical data explained by the source reconstruction. In the resolution analysis, we calculate measures based upon the resolution matrix (Hauk et al., 2011), which theoretically relates the activity based at dipole *i* to the estimate at dipole *j*.

**Figure 1:**
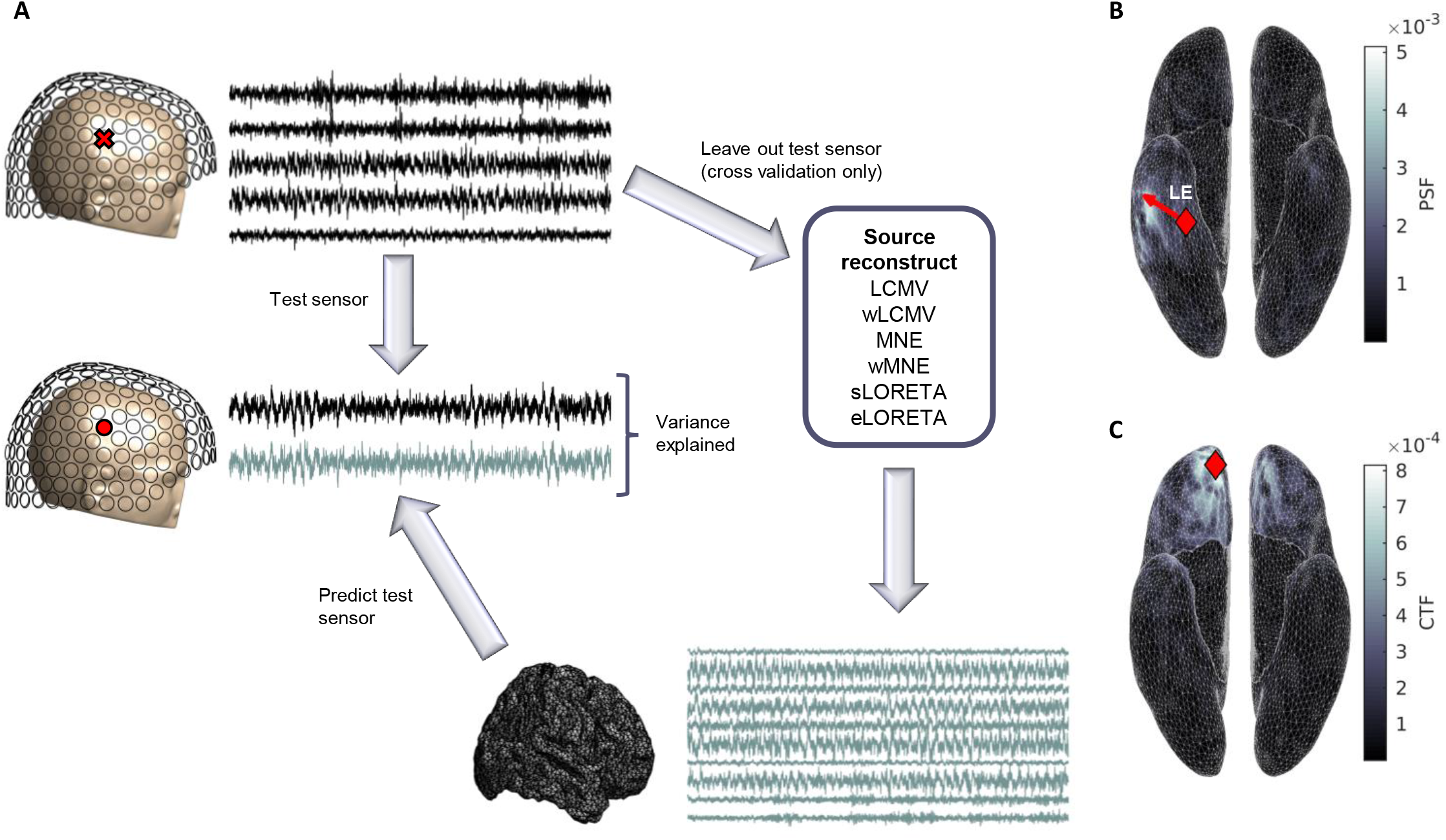
Methods of comparing source reconstruction algorithms. (**A**) Variance explained analysis. For the cross-validation procedure, the 274 channel MEG data is split into a single test sensor and the remaining 273 training sensors. For each algorithm, the training data is source reconstructed to gain an estimate of source dynamics. This source reconstructed activity is then forward mapped to the left out test sensor, and the variance of the data recorded at this sensor explained by the brain activity is calculated as squared correlation. This is repeated 274 times, using a different test sensor for each fold. Algorithms with high variance explained give a more realistic estimate of source activity. For the non-cross-validated procedure, all 274 sensors are source reconstructed, forward mapped, and predicted data for all sensors correlated with empirical data. (**B**) For a seed voxel, the corresponding column of the resolution matrix (the point spread function; PSF) relates unit ‘true’ activity at that voxel to source estimates at each voxel. The distance between the seed voxel and the voxel at the peak of the PSF is the localization error (LE), measured in mm. For demonstration purposes, we show the PSF of the wMNE solution for a seed voxel (red diamond) with maximum LE over all voxels and subjects. The arrow points from the seed to the peak of the PSF, and the length of the arrow is LE. (C) For a seed voxel, the corresponding row of the resolution matrix (the cross talk function; CTF) quantifies the influence of ‘true’ activity at all voxels on the source estimate of the seed voxel, i.e. it measures the amount of contamination of the source estimate from other voxels. The spatial extent of cross talk (SECT) is the CTF-weighted sum of distances to each voxel from the seed. For demonstration purposes, we show the CTF of the eLORETA solution for a seed voxel (red diamond) with maximum SECT over all voxels and subjects. In B-C, figures are plotted on the smoothed cortical surface for visualization purposes only.

#### 2.2.1 Variance explained analysis

A source reconstruction algorithm should accurately estimate the brain activity underpinning the M/EEG data. Therefore forward modelling to a new sensor should accurately predict the activity at that sensor. Here, we use a leave-one-out procedure to assess source reconstruction solutions based on this assumption. Data from *N_x_* – 1 sensors are source reconstructed to estimate brain activity ŝ (Equation 2). The predicted data at the remaining ‘test’ sensor is then estimated as 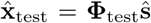, where **Φ**_test_ is the row of the leadfield matrix corresponding to the test sensor. This procedure is repeated for all the *N_x_* sensors. The variance of the empirical data explained by this predicted data, 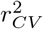, is quantified by the squared Pearson correlation coefficient between 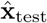 and the empirical data recorded by the test sensor, averaged over all test sensors, i.e.,

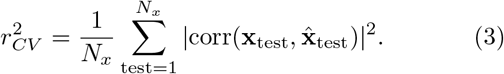

In a less stringent but more computationally efficient estimate, we skipped the leave-one-out procedure, reconstructing the source solution from all sensors, and in this case we calculate

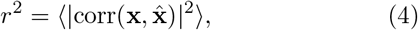

where 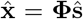. In particular, the ratio 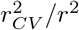 quantifies of the robustness of an algorithm to cross validation.

#### 2.2.2. Resolution analysis

We further used two measures derived from the resolution matrix (Hauk et al., 2011) to compare algorithms: the localization error and the spatial extent of cross talk.

Substituting Equation 1 into Equation 2, we obtain

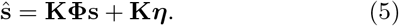

Although we cannot know the true source dynamics **s**, we know they are related to the estimated source dynamics ŝ by the equation

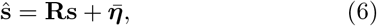

where 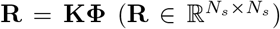 is the *resolution matrix* and 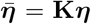 is a noise term (specifically, it is the source space projection of the measurement noise). The *i*’th column of **R** is called the point spread function of dipole *i* (PSFj), which corresponds to the influence of activity at dipole i on the estimate at all dipoles in the source reconstruction. The localization error of dipole i (LE¿) (Hauk et al., 2019) is thus given by

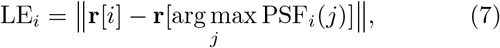

where **r**[*k*] is the location of dipole *k*. That is, a dipole i located at **r**[*i*] most strongly influences the estimate of dipole *j* at location **r**[*j*], and LE is the Euclidean distance between these two dipoles. The *i*’th row of **R** is called the cross talk function of dipole *i* (CTF_*i*_), which corresponds to the weights with which each dipole influences the estimate of activity at dipole *i*. We can therefore quantify leakage using a metric previously called ‘spatial dispersion’ (Hauk et al., 2019), but here called the spatial extent of cross talk (SECT) to avoid misinterpretation (see section 4.3 for a justification of this difference in naming convention). The SECT influencing dipole *i* is calculated as

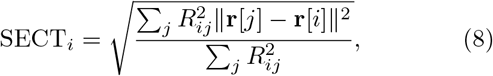

where *R_ij_* is element (*i*, *j*) of **R**. In the idealized solution with no leakage, **R** = **I**. As non-diagonals become nonzero, so does SECT.

#### 2.2.3. Comparing source reconstruction algorithms using parcellated data

We extended the comparisons between source reconstruction algorithms to parcellated data. For the variance explained analysis, cortical parcellation was performed by taking the first principal component of all voxels within a ROI (Tait et al., 2019) to obtain a single time course for the ROI. The representative time course of the ROI was then mapped back to voxel space using the column of the PCA mixing matrix corresponding to the first principal component. The variance explained analysis with cross-validation was then performed as described in section 2.2.1. For the resolution analysis, we defined frac-tional localization error for ROI *ω* (fLE_*ω*_) as the fraction of dipoles within ω that had peaks of the PSF outside of ω. As a parcellated counterpart to SECT, we calculated the zero-lag correlation between time courses of anatomically neighbouring ROIs, since highly correlated neighbouring regions suggests large spatial leakage. This was averaged over all pairs of anatomical neighbours, to obtain the mean neighbour correlation (mNC).

#### 2.2.4. Statistical analysis

For all measures, values reported were averaged over all dipoles or ROIs unless otherwise stated. Statistical analysis of metrics across algorithms required tests accounting for repeated measures to address the within-subject experimental design. Friedman tests were used for group level analyses, and pairwise comparisons were performed using the Wilcoxon signed rank test. All *p*-values reported for pairwise comparisons were false discovery rate corrected using the Benjamini-Hochberg procedure.

### 2.3. Reducing the HCP-MMP atlas

We proposed a data-driven approach to reduce the HCP-MMP atlas from 360 to 250 cortical ROIs. This number was chosen such that the number of ROIs was less than or equal to the rank of our preprocessed MEG data. This was motivated by the fact it is commonplace to orthogonalize parcellated data, e.g. for functional connectivity analyses, and therefore by using fewer ROIs than the rank of the data we avoid rank deficiency.

Glasser et al. (2016) grouped 180 cortical ROIs per hemisphere into 22 anatomical ‘clusters’ of regions, and our reduced atlas aimed to maintain these clusters. Consider a cluster Ω. For a given ROI *ω* ∈ Ω, the influence of that ROI over the MEG was defined as

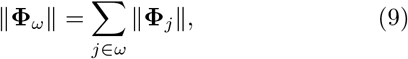

where Φ_*j*_ is the column of the leadfield matrix corresponding to dipole *j*. Here, we take the average value of ||Φ_*ω*_ || over subjects using our MRI derived leadfields. This average is demonstrated to be representative of all participants in Figure S1. Then the corresponding influence of Ω is calculated as

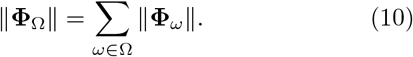

The number of ROIs in Ω should ideally be proportional to ||Φ_Ω_||. Then, given a target number of ROIs per hemisphere (here chosen to be 125), a simple algorithm was used to determine the optimal number of ROIs for each cluster, such that the target total is achieved. The algorithm is described in detail in Appendix B. Subsequently, we used the detailed neuroanatomical results presented by Glasser et al. (2016) to merge neigbouring ROIs within a cluster that had similar neuroanatomical attributes (prioritising resting-state functional connectivity) until the cluster had the optimal number of ROIs and a uniform distribution of ROI strengths. A full list of ROIs and descriptions of which ROIs were merged is given in Supplementary Text S1.

### 2.4. Participants

Eleven healthy participants (8 female, 3 male) were recruited from Cardiff University School of Psychology participant panel (age range 19-24 years, mean age 20.45 years). All participants had normal or corrected-to-normal vision, and none reported a history of neurological or psychiatric illness. Written consent was obtained from all participants. The study was approved by the Cardiff University School of Psychology Research Ethics Committee.

### 2.5. MEG and MRI data acquisition

Whole-head resting-state MEG recordings were made using a 275-channel CTF radial gradiometer system (CTF Systems, Canada) at a sampling rate of 1200 Hz. An additional 29 reference channels were recorded for noise cancellation purposes and the primary sensors were analysed as synthetic third-order gradiometers (Vrba and Robinson, 2001). One sensor was turned off during recording due to excessive sensor noise (i.e., *N_x_* = 274 gradiometers). Subjects were instructed to sit comfortably in the MEG chair while their head was supported with a chin rest and with eyes open focus on a fixation point on a grey background. Horizontal and vertical electro-oculograms (EOG) were recorded to monitor blinks and eye movements. The horizontal electrodes were placed on temples, and vertical ones, above and below the eye. For MEG/MRI coregistration, the head shape with the position of the coils was digitised using a Polhemus FASTRAK (Colchester, Vermont). Each recording session lasted approximately eight minutes.

All participants also underwent a whole-brain MRI scan on a Siemens 3T Connectom MRI scanner and a 32-channel receiver head coil (Siemens Medical Systems). We used a T1-weighted magnetization prepared rapid gradient echo sequence (MPRAGE; echo time: 3.06 ms; repetition time: 2250 ms sequence, flip angle: 9°, field-of-view: = 256 × 256 mm, acquisition matrix: 256 × 256, voxel size: 1 × 1 × 1 mm).

### 2.6. Data preprocessing and forward modelling

Continuous raw MEG data was imported to Fieldtrip (Oostenveld et al., 2011), bandpass filtered at 1-100 Hz (4th order two-pass Butterworth filter), with the exception of the 100+ Hz signal in section 3.2.2, for which a 1 Hz highpass filter was applied instead during preprocessing. Data was subsequently notch filtered at 50 and 100 Hz to remove line noise. Visual and cardiac artifacts were removed using ICA decomposition, using the ‘fastica’ algorithm (Hyvärinen, 1999). Identification of visual artifacts was aided by simultaneous EOG recordings. Between 2 and 5 components were removed for each subject. Data was then downsampled to 256 Hz and a randomly selected 30 second epoch was chosen. The Fieldtrip functions ‘ft_artifact_jump’ and ‘ft_artifact_clip’ were used to validate that this epoch was free from clip and jump artifacts.

From T1-weighted MRI image, extraction of the inner skull, scalp, pial, and grey matter/white matter boundary surfaces was performed with the Freesurfer (Dale et al. (1999), http://surfer.nmr.mgh.harvard.edu). These surfaces and the MRI scan were imported into the Brainstorm software (Tadel et al., 2011) and an automated procedure used to align these data to the MNI coordinate system. The midpoint between the pial surface and grey matter/white matter boundary was extracted and downsampled to 10,000 homogeneously spaced vertices (median distance between vertices 4.32 mm; Figure S2) to generate a cortical surface of dipole locations using the ‘iso2mesh’ software (Fang and Boas, 2009) implemented in Brainstorm. The inner skull surface was similarly downsampled to 500 vertices. These surfaces were then exported to Matlab, where the scalp surface was used to align the structural data with the MEG digitizers. The aligned MEG gradiometers, inner skull surface, and cortical surface were then used to construct a realistic, subject specific, single shell forward model (Nolte, 2003). Unless stated otherwise, dipole orientations were fixed normal to the cortical surface (Dale et al., 2000; Hillebrand and Barnes, 2003) under the assumption that M/EEG signals are primarily generated by the postsynaptic currents in the dendrites of large vertically oriented pyramidal neurons in layers III, V, and VI of the cortex (Olejniczak, 2006).

For the parcellation analysis, the cortical surface was further aligned to the Human Connectome Project Multimodal Parcelation (HCP-MMP) atlas (179 cortical ROIs per hemisphere, excluding the hippocampus; Glasser et al. (2016)) in Freesurfer, and Matlab was used to merge ROIs according to the parcellation reduction procedure of section 2.3.

## 3. Results

### 3.1. Comparing source reconstruction algorithms

We began by comparing six source reconstruction algorithms in broadband (1-100 Hz) data. Figure 2 shows the correlation between the source space solution of pairs of algorithms, and hierarchical clustering of the algorithms based on these correlations. MNE and wMNE were mostly independent of other algorithms (with the exception of a small positive correlation between eLORETA and wMNE), while LCMV, wLCMV, sLORETA, and eLORETA formed a cluster with correlations greater than ~ 0.3. Most strongly correlated in this cluster were sLORETA and wLCMV, which had correlations close to one. Appendix C provides an analytical proof that the wLCMV and sLORETA solutions have similar but non-identical forms, potentially explaining this result. eLORETA was the least strongly correlated with this cluster, and the first to be cut from the cluster using the iterative clustering algorithm. This result, combined with the presence of a small positive correlation with wMNE, suggests that the eLORETA solution is an intermediate between the solutions of the wMNE algorithm and the LCMV/wLCMV/sLORETA cluster.

**Figure 2:**
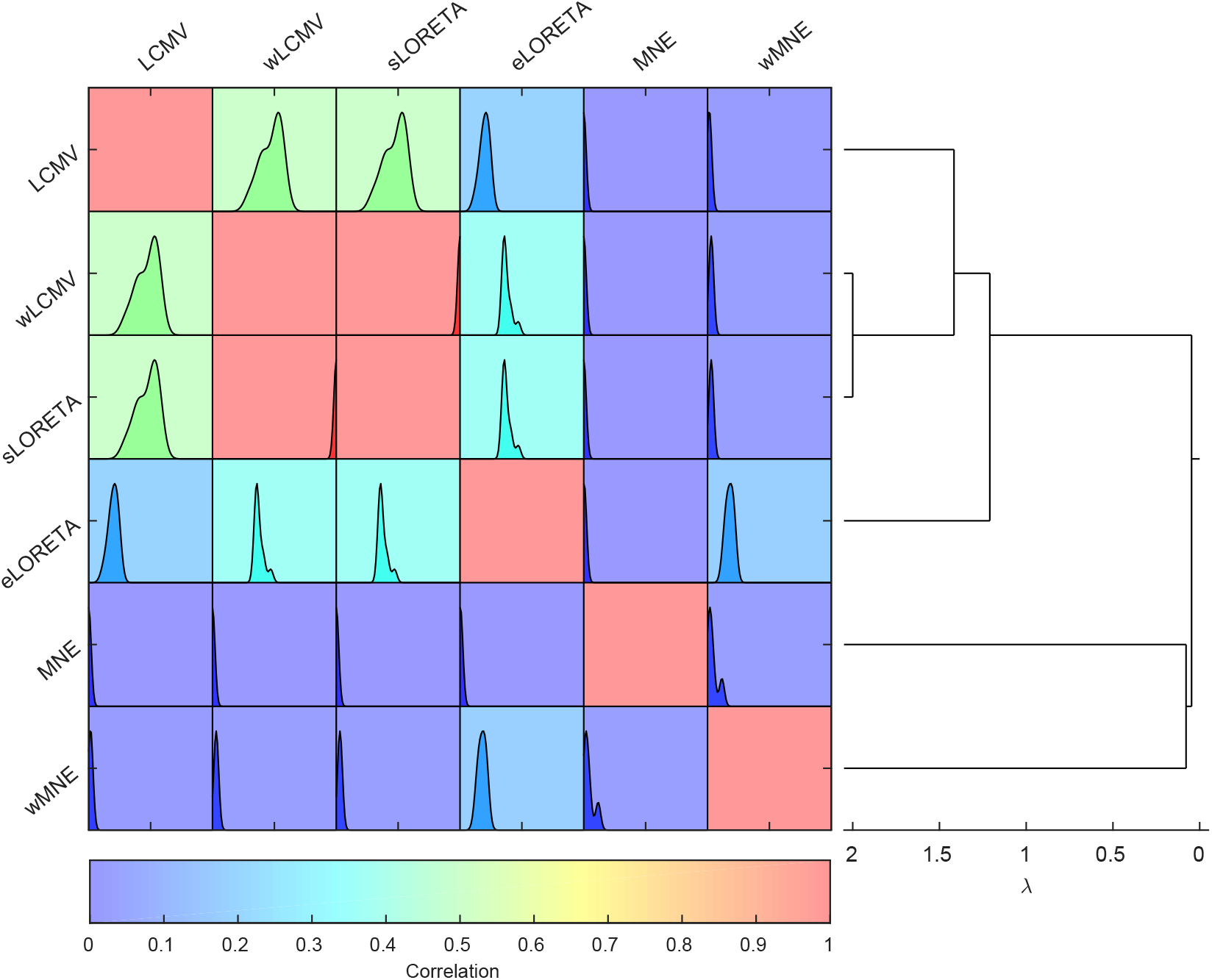
Correlation between different source reconstruction algorithms.

Each algorithm was used to source reconstruct the broadband data, and correlation in the source space solutions of each pair of algorithms was calculated. The correlation matrix is shown on the left. For each pair of algorithms, the background colour represents median value of correlation over all participants, whilst the inlaid plot shows the distribution over all subjects (the x-axis is calibrated evenly for all plots, in the range [0,1]). No negative correlations were found. Hierarchical clustering of the median network is shown on the right to group algorithms based on similarity. Clustering used a spectral normalized cuts algorithm (Shi and Malik, 2000), and the x-axis shows algebraic connectivity (λ; i.e. cost of the cut). Therefore clusters with higher λ are more strongly clustered, i.e. contain higher correlations between algorithms.

### 3.1.1. Variance explained analysis

Performance of the algorithms was quantified using an analysis of variance of empirical data explained by the source reconstruction, 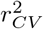 (see section 2.2.1 and Equation 3). The values of 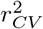, averaged over sensors, are shown for each algorithm in Figure 3A, and their spatial extent are shown in Figure S3. There was a significant effect of algorithm on 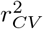 (χ^2^ = 55, *p* = 1.31 × 10^−10^, Friedman test), and pairwise comparisons demonstrated significant differences between all pairs of algorithms (*p* = 9.77 × 10^−4^ for all pairwise tests).

**Figure 3:**
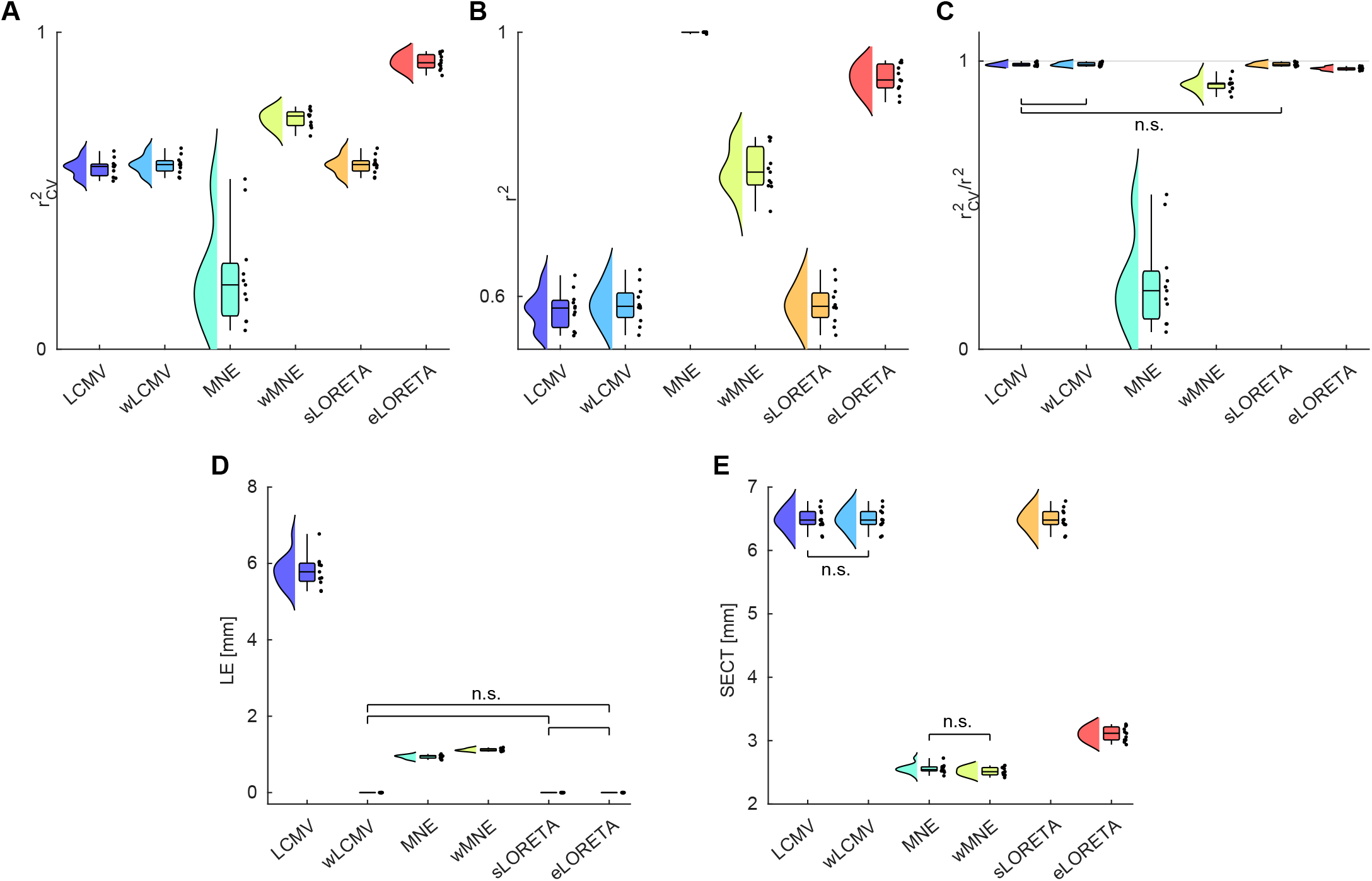
Statistics from variance explained and resolution analysis. (A) Cross-validated variance of empirical data explained 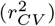 for each source reconstruction algorithm. (B) Variance of empirical data explained without cross-validation (*r*^2^). (C) Robustness to crossvalidation, quantified by the ratio 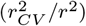. Values close to 1 (marked by a horizontal grey line) demonstrate that the source reconstruction procedure is robust and not overfitting to the data. (D) Localization error (LE). (E) Spatial extent of cross talk (SECT). For all figures, group-wise analyses demonstrated a significant effect of algorithms. Pairwise analyses identified significant differences for all pairs except those marked as non-significant (n.s.), following false discovery rate correction. In the box plots, the range (whiskers), interquartile range (boxes), median (horizontal line), are shown, with values for each participant marked by dots to the left.

Of the algorithms, eLORETA had the largest 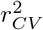, explaining 90.6 ± 0.7% of the data (mean ± SEM), followed by wMNE (72.6 ± 0.9%). Interestingly, sLORETA and wLCMV demonstrated similar 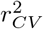 (sLORETA: 58.144 ± 0.862%; wLCMV: 58.141 ± 0.861%), although wLCMV performed slightly and consistently better than sLORETA in all subjects (*p* = 9.77 × 10^−4^, Wilcoxon signed-rank test). LCMV had marginally lower 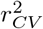 than sLORETA and wLCMV (57.3 ± 0.9%). All algorithms except MNE were robust to cross validation, with similar values of 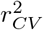 and *r*^2^ (i.e., variance explained without cross-validation, see section 2.2.1 and Equation 4) as shown in Figure 3B-C. Interestingly, while the MNE algorithm performed poorly in the cross-validated analysis 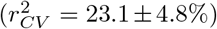, it had *r*^2^ = 1 for all participants, suggesting that MNE has a strong tendency to over-fit to the data.

### 3.1.2. Resolution analysis

In the first resolution analysis, we quantified the localization error (LE) for each algorithm (see section 2.2.2 and Equation 7). There was a significant effect of algorithm on LE (Figure 3D and Figure 4A, χ^2^ = 55, *p* = 1.31 × 10^−10^), with significant differences between all pairs of algorithms (*p* = 0. 0012 for all pairwise tests) except for comparisons between wLCMV, sLORETA, and eLORETA (*p* = 1). As previously reported numerically (Pascual-Marqui et al., 2018; Hauk et al., 2019) and theoretically expected (Pascual-Marqui, 2007), eLORETA and sLORETA had zero localization error. Interestingly, wLCMV also had zero LE in our results. In Appendix C, we provide mathematical proof that, theoretically, one might expect non-zero LE in the wLCMV beamformer, and small nonzero LE may be found if a much finer grid is used for source reconstruction. Nevertheless, under the grid spacing used in the current study (Figure S2), which is commonly used in practice, the LE of the wLCMV solution is numerically negligible. MNE had the next lowest LE (0.94±0.01 mm), followed by wMNE (1.12 ± 0.01 mm), whilst LCMV had the largest LE (5.8 ± 0.13 mm).

**Figure 4:**
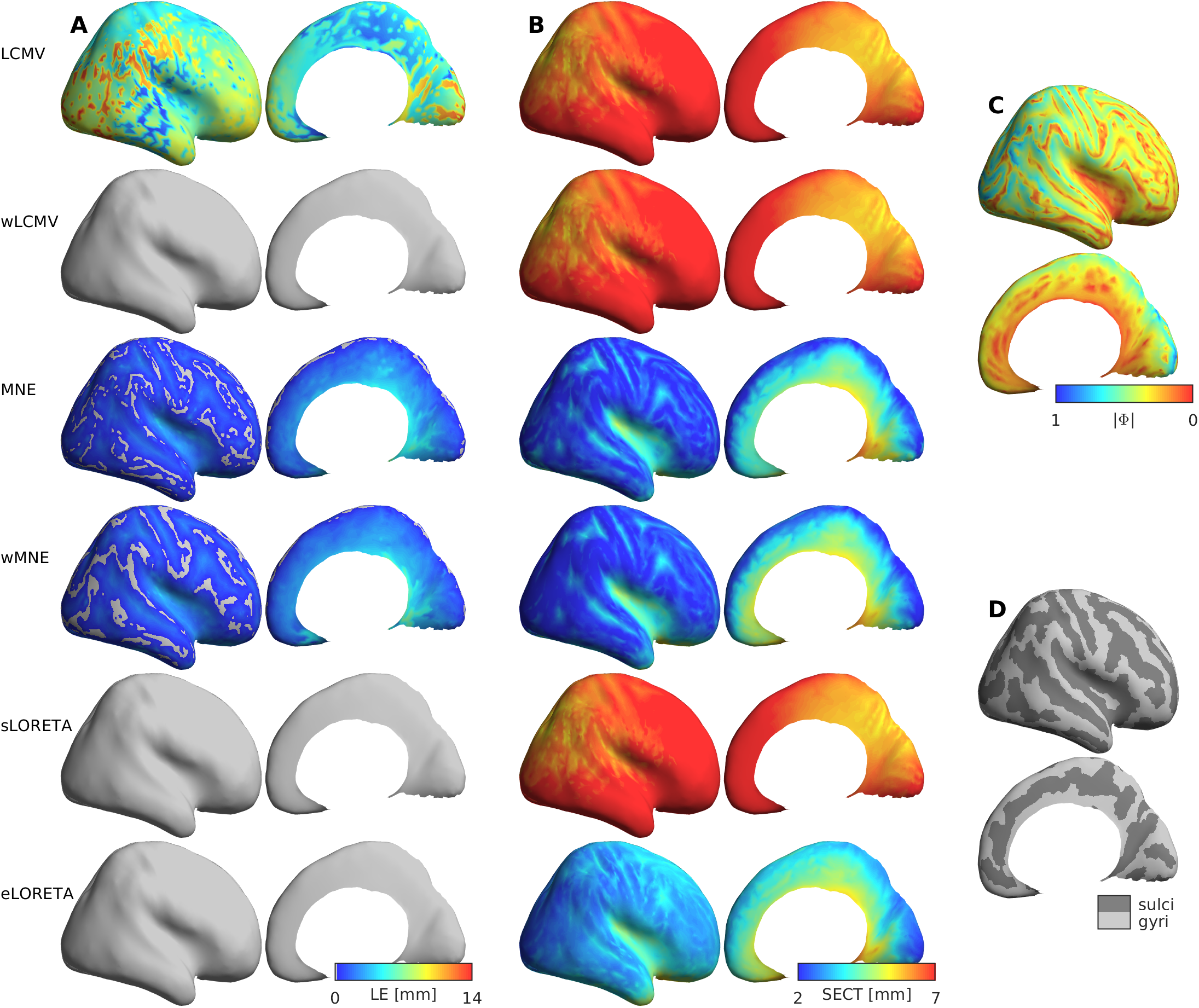
Spatial distributions of resolution metrics.

(A-B) Lateral and medial views (left and right columns) of LE and SECT (respectively) for each dipole in each algorithm. (C) The norm of the leadfield for each dipole. Note that in C, the colour bar is flipped with respect to A-B, so bright areas have low leadfield norm (weak sources), whilst dark areas are strong. (D) Anatomical maps of gyri and sulci, derived in Freesurfer. For all figures, distributions are shown on a smoothed cortical surface for visualization purposes, and are shown for the same representative participant.

The second resolution analysis compared the spatial extent of cross talk (SECT) influencing the source estimates (see section 2.2.2 and Equation 8). There was a significant effect of algorithm on SECT (Figure 3E and Figure 4B, χ^2^ = 53.50, *p* = 2.65 × 10^−10^), with significant differences between all pairs of algorithms (*p* < 0.0011 for all pairwise tests) except for LCMV vs wLCMV (*p* = 1) and MNE vs wMNE (*p* = 0.3913). wMNE had the lowest SECT (2.52 ± 0.02 mm), followed by MNE (2.56 ± 0.02 mm), and eLORETA (3.11 ± 0.03 mm). LCMV and wLCMV had identical values of SECT (6.49±0.05 mm), since setting columns of the spatial filter to unit norm has no effect on SECT. The sLORETA algorithm had values similar to wLCMV/LCMV (6.48 ± 0.05 mm), but for all subjects had of the order 10^−3^ mm lower SECT than the beamforming algorithms.

The results above were obtained from forward models with anatomical dipole orientations that are normal to the cortical surface (Figure S4). In supplementary Figure S5, we showed that the relative ranking of algorithms is maintained in most measures (i.e., 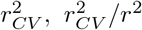, LE and SECT) if using eigendecomposition to define dipole orientation (Sekihara et al. (2004); Figure S4), suggesting that our results do not depend on anatomical orientation constraints.

### 3.2. Effects of signal-to-noise ratio to source reconstruction algorithms

A key factor affecting the source reconstruction is signal-to-noise ratio (SNR). In resting-state data, where there is no good model of ‘brain noise’, the predominant sources of noise to be considered are noise from movement artifacts as well as extrinsic or measurement noise including quantisation errors, ambient magnetic fields, etc (Parkkonen, 2010). A range of factors can increase the SNR of the resting-state MEG, including the use of active noise cancellation systems, reference sensors, signal space projection, and bandpass filtering (Parkkonen, 2010). Since resting-state data is not time-locked to a stimulus, this noise cannot be readily reduced by averaging over epochs. This section considered how source reconstruction algorithms are affected by different levels of external white noise and by intrinsic structured noise at different frequency bands.

#### 3.2.1. Effects of external white noise

For each participant, each algorithm, and each MEG sensor, we added i.i.d. Gaussian white noise with different amplitudes (i.e., a standard deviation of *σ* multiplied by the average standard deviation of the data) to sensor-level MEG data, which resulted in a range of SNRs. Here, we treat the original empirical sensor-level MEG data as ‘signal’ and the added white noise as ‘noise’. Due to the high computational cost of cross validation across participants, algorithms and noise levels, we quantify the effects of SNR on source reconstruction performance with the variance explained measure *r*^2^ (Equation 4) instead of 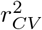. When the level of noise *σ* increases (i.e., lowering SNR), an ideal source reconstruction would only explain the signal but not the (artificially added) noise, with *r*^2^ decreasing as *σ* increases, whilst the source space solution remains unaltered. Therefore, in Figure 5A, which shows the variance explained measure *r*^2^ with a noise level *σ* normalized by that with *σ* = 0, the ideal solution should follow the theoretical performance 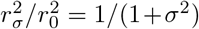 (dotted black line). In Figure 5B, which shows the Pearson correlation coefficient between source reconstruction of MEG data with and without added noise, *r*(*s*, *s*_0_), the ideal solution should remain at one for all *σ*.

**Figure 5:**
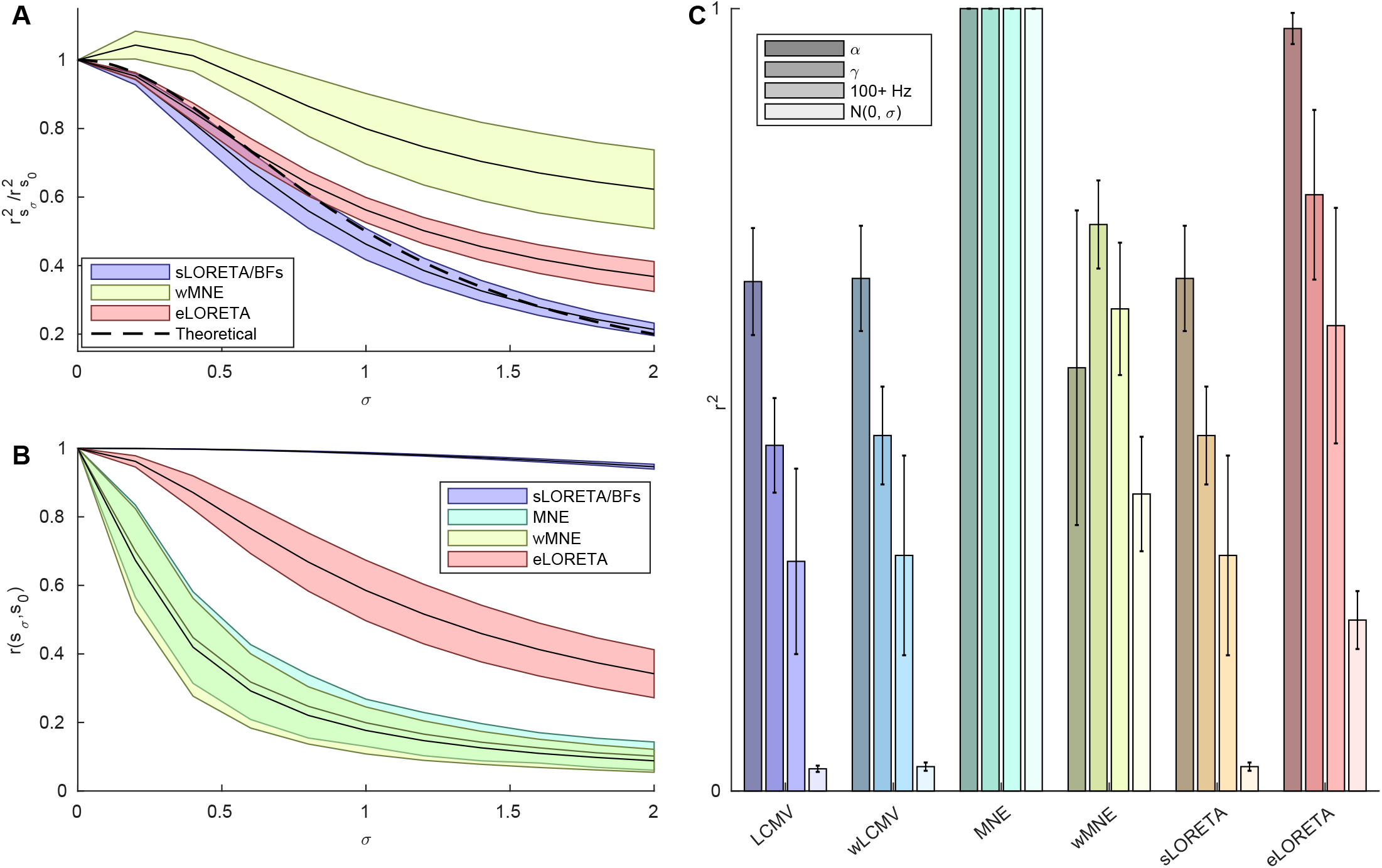
Effects of varying signal-to-noise ratio.

(A-B) SNR was varied by adding white noise to the MEG data with a standard deviation of *σ* times the standard deviation of the data. Variance explained by the source reconstruction of the MEG data (A) and correlation between the signal in source space in the case with added noise and no added noise (B) were calculated for each SNR. In A, values are normalized by values when *σ* = 0, and the variance explained by a theoretically ideal source reconstruction (see section 3.2.1 for description) is shown by a dotted black line. sLORETA and the LCMV/wLCMV beamformers give near identical solutions, so are plotted as a single colour (sLORETA/BFs). MNE had a value of one for all participants and all values of *σ*, so is not included in the plot. Lines are the median, whilst shaded regions are interquartile range. (C) SNR was varied by bandpass filtering the data into different frequency bands. Bands are sorted by decreasing hypothesised SNR. The alpha (8-13 Hz) band is hypothesised to have high SNR as it is predominantly high amplitude oscillations. The gamma (30-100 Hz) band is hypothesised to be a mixture of low amplitude gamma oscillations and noise, so has a lower SNR. The 100+ Hz band is hypothesised to contain very little signal, being predominantly noise. Finally, pure white noise with no added signal was source reconstructed, giving an SNR of 0.

MNE always explains the full data including the noise (*r*^2^ = 1 for all values of *σ*), whilst *r*(*s*,*s*_0_) rapidly decreases with larger *σ*. This supports the hypothesis that MNE overfits to the data (i.e., Figure 3C). For wMNE, *r*^2^ increases at low levels of noise (*σ* <~ 0.5), and decreases at higher *σ* values, but it is always much greater than the theoretical variance explained for an ideal solution. Furthermore, similar to MNE, the *r*(*s*, *s*_0_) for wMNE decreases sharply as *σ* increases. These results suggest that wMNE is also strongly affected by the presence of external noise. sLORETA and the LCMV/wLCMV beamformers give almost identical solutions in both analyses, with *r*^2^ consistently decreasing as *σ* is increased and maintaining high *r*(*s*, *s*_0_) (> 0.936 for all *σ*), suggesting these solutions are largely unaffected by the addition of white noise. eLORETA also demonstrated consistent decreases in the *r*^2^ measure as levels of noise were increased, but deviated from the theoretical value particularly for high values of *σ*. Additionally, the *r*(*s*, *s*_0_) of eLORETA dropped more rapidly with increasing *σ* than for sLORETA and the beamformers, suggesting that eLORETA-based source solutions would be altered under varying SNR.

#### 3.2.2. Effects of empirical noise across frequency bands

To consider the effects of SNR with realistic structured noise, we subsequently altered the SNR by bandpass filtering the data, capitalizing on previous findings that noise in human electrophysiological data, especially those originated from muscle artifacts, are frequency-dependant (Criswell, 2010). In contrast to the previous section, no noise was added artificially. We band-pass filtered the sensor-level MEG data into three frequency bands: the alpha band (8-13 Hz), the gamma band (30-100 Hz) and a high frequency band over 100 Hz, to compare the variance explained measure *r*^2^ between source reconstruction algorithms across frequency bands. The alpha band data has been shown to contain primarily oscillations of neuronal origin and is less affected by muscle artifacts (O’Donnell et al., 1974; Goncharova et al., 2003), meaning that it has a relatively high hypothetical SNR. The gamma (30-100 Hz) band data is a mixture of low amplitude neuronal oscillations and strong muscular contamination (Kumar et al., 2003; Whitham et al., 2007, 2008), including common artifacts from extraocular muscles (Yuval-Greenberg et al., 2008; Carl et al., 2012), resulting in a medium hypothetical SNR. MEG data at the 100+ Hz band is likely to be dominated by noise, because this band is close to the upper range limit of neuronal oscillations to be recorded by MEG sensors (Parkkonen, 2010), resulting in a low hypothetical SNR. Furthermore, to provide a baseline for comparison, we additionally replaced empirical MEG recordings with time sources of pure white noise for source reconstruction, which effectively leads to a zero SNR condition.

Figure 5C shows the results of frequency-specific *r*^2^. sLORETA, the wLCMV/LCMV beamformers, and eLORETA, all explain less variance as the hypothetical SNR decreases from the alpha band to the zero SNR condition, suggesting all these algorithms are sensitive to and affected by realistically structure noise. Consistent with the results from broadband data (Figure 5A), MNE has *r*^2^ values close to 1 in all conditions, suggesting that MNE overfits to the data regardless the level of noise.

#### 3.3. The reduced HCP-MMP atlas

We proposed a data-driven method (see section 2.3) to reduce the 360 ROIs in the HCP-MMP atlas to 250 ROIs, fewer than the rank of our MEG data and hence appropriate for further analysis (Colclough et al., 2015). The essential criterion for ROI reduction is to merge neighbouring ROIs that have small influences over MEG sensor-level data (given by the MEG leadfield matrix, see Equation 10), while maintaining all the clusters in the original HCP-MMP atlas with similar microstructural, functional and connectivity profiles (Glasser et al., 2016). As such, the reduced HCP-MMP atlas is optimized for MEG source reconstruction.

The reduced atlas is shown in Figure 6, and a detailed description of all ROI reductions to the HCP-MMP atlas is given in Supplementary Text S1. Merged ROIs were found mostly in the less superficial areas such as medial temporal regions (including the ventral visual stream and medial temporal cortex), the lateral sulcus (including early auditory, insular and opercular cortices), and medial regions (including the cingulate, medial prefrontal, and orbitofrontal cortices) (Figure 6C and F). By merging ROIs in these regions that have small influences over the MEG, the distributions of influence of ROIs became more homogeneous (Figure 6A and D). For example, the insular cortex in the original atlas consisted of thirteen small and deep ROIs per hemisphere which had little influence on the MEG (Figure 6A–C). Our algorithm determined the optimum number of regions was four, so by merging insular ROIs which were anatomical neighbours and had similar resting-state functional connectivity as reported by Glasser et al. (2016), we achieved four larger ROIs with comparable influence over the MEG to ROIs in more superficial regions (Figure 6D–F). These merges can be justified by our resolution analysis (Figure 4B), which demonstrates that the insular had much lower resolution (i.e., higher SECT) than superficial regions for all source reconstruction algorithms.

**Figure 6:**
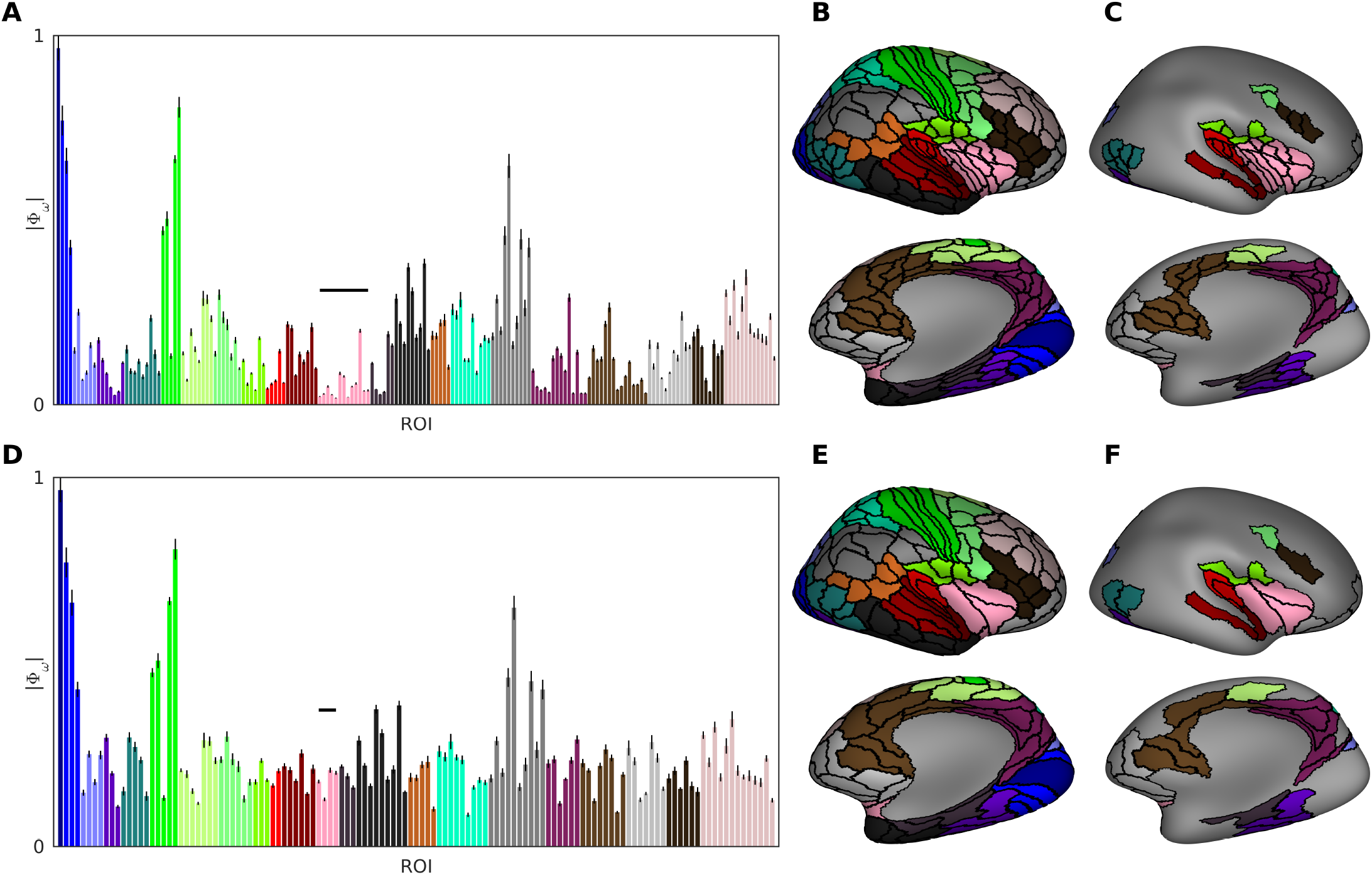
The reduced HCP-MMP atlas. (A-C) The original atlas (Glasser et al., 2016), consisting of 180 ROIs per hemisphere split into 22 larger ‘clusters’. (A) The norm of the leadfield, summed over all voxels in an ROI *ω*, quantifying the strength with which *ω* influences the MEG. Clusters are marked by different colours, whilst each bar is an individual ROI. (B) Each ROI’s anatomical location, plotted on the inflated Freesurfer average brain. Top shows the lateral surface, whilst the bottom is angled to show the medial and ventral surfaces. (C) As in B, but plotting only ROIs which are modified in the reduced atlas to highlight changes. (D-F) The reduced atlas, consisting of 125 ROIs per hemisphere, plotted similarly as in A-C respectively. We reduce the atlas by stating that each cluster should have a number of ROIs proportional to its influence on the MEG (A and D). For example, cluster 12 (the insula, the pink cluster marked in A and D by a black bar) has very many weak ROIs, so we combine these ROIs into fewer, larger, more influential ROIs. The result of this is a more uniform distribution of ROI strengths. Whilst the calculation of numbers of ROIs per cluster was automated, the choice of which specific ROIs to be combined was made by hand from studying anatomical and functional closeness, based on the study of Glasser et al. (2016).

#### 3.4. Comparing source reconstruction algorithms using parcellated data

Based on the reduced HCP-MMP atlas Figure 6E, we parcellated all voxelwise solutions of source reconstruction algorithms into 250 ROIs. For each ROI, its representative time course was obtained from the first principal component across all voxels within that ROI. We then compared the source reconstruction algorithms using the parcellated, representative time courses of ROIs. The variance explained analysis and resolution metrics were adapted accordingly for parcellated data (see section 2.2.3).

The values of cross-validated variance explained 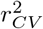 are shown for each algorithm in Figure 7A. There was a significant effect of algorithm on 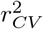 (χ^2^ = 49.4, *p* = 1.84 × 10^−9^, Friedman test), with significant pairwise differences between all pairs of algorithms (*p* = 0.0056 for wLCMV vs sLORETA, *p* = 0.0012 otherwise) except LCMV vs wLCMV (*p* = 0.7646) and LCMV vs sLORETA (*p* = 0.7646). Compared with results from voxelwise data (Figure 3A), atlas-based parcellation resulted in lower 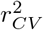 values. This is expected, because the PCA-based representative time course do not contain all the information in an ROI. Nevertheless, eLORETA still had the largest 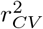, explaining 77.0±1.1% of the data, followed by wLCMV (48.4±0.8%), sLORETA (48.4 ± 0.8%) and LCMV (48.3 ± 0.7%). The MNE and wMNE solutions performed poorly when parcellation was applied to the data, explaining 0.14 ± 0.04% and 4.55 ± 0.48% of the data, respectively. Whilst MNE was previously reported to perform poorly for unparcellated data (section 3.1), this result highlighted that neither MNE or wMNE provide reliable solutions for parcellated data.

**Figure 7:**
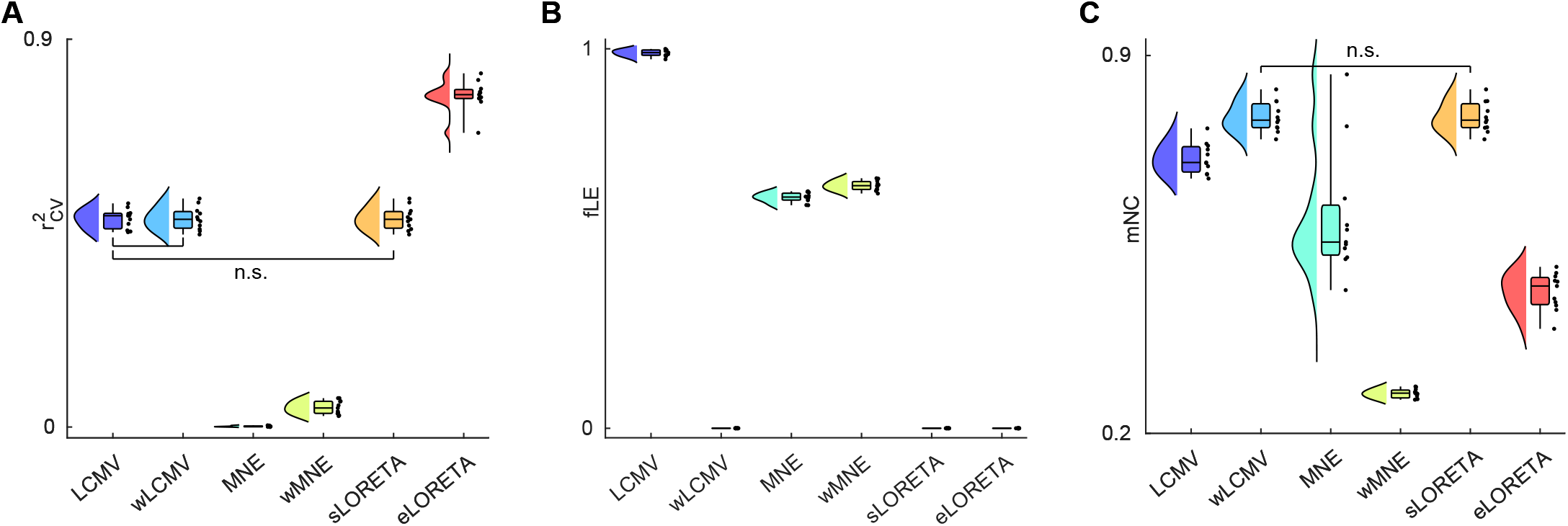
Parcellation based statistics.

(A) Variance explained in sensor space by the parcellated dynamics. (B) Fractional localization error (fLE). (C) Mean neighbour correlation (mNC) of ROI time series. For all figures, group-wise analyses demonstrated a significant effect of algorithms. Pairwise analyses identified significant differences for all pairs except those marked as non-significant (n.s.), following false discovery rate correction. Values for whiskers, boxes, etc. are described in Figure 3.

Fractional localization error (fLE) was calculated as a parcellation based counterpart to LE. By definition, zero LE results in zero fLE, so consistent with unparcellated results we found zero fLE for sLORETA, wLCMV, and eLORETA. There was a significant effect of algorithm on fLE (χ^2^ = 55, *p* = 1.31 × 10^−10^), with significant differences in all tests (*p* = 0.0012 for all comparisons) except those between algorithms with zero fLE (*p* = 1). Consistent with the LE in unparcellated results (Figure 3D), MNE had the lowest non-zero fLE (0.608 ± 0.004), followed by wMNE (0.639 ± 0.004). LCMV had very high fLE (0.989±0.003), suggesting that the majority of source reconstructed signal is attributed to a wrong ROI.

Finally, mean neighbour correlation (mNC) was used as a measure of resolution/leakage of the parcellated data, a counterpart of the SECT resolution measure for voxelwise analyses. Low mNC is suggestive of low leakage, since resolution is high enough to distinguish activity at neighbouring ROIs. There was a significant effect of algorithm on mNC (χ^2^ = 47, *p* = 5.68 × 10^−9^), with significant differences between all algorithms (*p* = 0.0450 for LCMV vs MNE, *p* ≤ 0.0056 otherwise), except wLCMV vs sLORETA (*p* = 0.7002). wMNE had the lowest mNC (0.273 ± 0.003), followed by eLORETA (0.4622 ±0.011), MNE (0.599 ±0.035), LCMV (0.710 ± 0.009), sLORETA/wLCMV (0.786 ± 0.008, differences of the order ±10^−5^ to ±10^<4^). Although there is no direct dependency between mNC and SECT, our results suggested that ranking of the algorithms based on mNC is largely in line with the SECT results for unparcellated data.

## 4. Discussion

The current study consisted of two main aims, unified by the goal of developing a methodology to guide future source reconstruction analyses of resting-state MEG. First, we systemically assessed which of the many inverse algorithms for source reconstructing MEG data was most suitable for use with empirical resting-state data. This was achieved through the measures of variance explained and resolution metrics. Second, we presented a reduced atlas that is based on the high resolution, multi-modal parcellation of the Human Connectome Project (Glasser et al., 2016), which is optimized for use with resting-state MEG data. Main contributions and implications of our results are discussed in the following sections.

### 4.1. Comparison of source localization algorithms

Our comparisons and recommendations of source localization algorithms are summarized in Table 1. Generally speaking, we recommend eLORETA as a first choice of algorithm for distributed cortical source reconstruction of resting-state MEG. In a wide range of conditions (broad-band/narrow band data, anatomical/eigendecomposition orientation constraints, unparcellated/parcellated), eLORETA (Pascual-Marqui, 2007, 2009) was the algorithm which could most accurately predict the MEG at the sensor level, explaining approximately 90% of variance in a cross-validation procedure. This suggests that the cortical dynamics estimated by eLORETA are representative of the macroscopic neural dynamics generating the measured MEG. Furthermore, in line with theoretical results (Pascual-Marqui, 2007) and simulated point dipoles (Pascual-Marqui et al., 2018), we found that eLORETA had zero LE and relatively lower leakage than LCMV, wLCMV and sLORETA. The fact these results were consistent across a number of conditions indicates that eLORETA is appropriate as a general tool for source reconstruction of resting-state MEG data. Using different methodologies to the one presented here, recent studies have also demonstrated that eLORETA outperformed other algorithms such as MNE and LCMV in resting-state EEG data (Liu et al., 2018; Finger et al., 2016), further supporting the use of this algorithm for resting-state source reconstruction.

**Table 1:** Summary of results and recommended usage of algorithms.

| Algorithm | Strengths | Weaknesses | Recommended use |
| --- | --- | --- | --- |
| LCMV | Uninfluenced by sensor noise. | High LE and SECT.<br>Nearly always misplaces sources during parcellation. | Never: High LE makes estimates unreliable. |
| wLCMV | Near-zero localization error.<br>Uninfluenced by sensor noise. | Poor ability to explain the data.<br>High cross talk, particularly after parcellation. | If low SNR/high sensor noise. |
| MNE |  | Highly non-robust to cross validation.<br>Highly overfits sensor noise. | Never: overfits data/noise. |
| wMNE | Reasonable ability to explain the data.<br>Low cross talk, including after parcellation. | Has small localization errors.<br>Strongly influenced by sensor noise.<br>Unreliable for narrow band data.<br>Misplaces sources after parcellation.<br>Poorly explains the data after parcellation. | If high resolution is a priority. |
| sLORETA | Zero localization error.<br>Uninfluenced by sensor noise. | Poor ability to explain the data.<br>High cross talk, particularly after parcellation. | If low SNR/high sensor noise. |
| eLORETA | Good ability to explain the data.<br>Zero localization error.<br>Reasonably low cross talk.<br>All of these properties maintained following parcellation. | Small amount of cross talk.<br>Reasonably influenced by sensor noise. | **First choice**, especially if data is to be parcellated. |

However, eLORETA is not a perfect solution. Whilst leakage is relatively low in eLORETA compared to LCMV, wLCMV, or sLORETA, other algorithms achieved better SECT (i.e., MNE and wMNE, Figure 3E) and mNC (i.e., wMNE, Figure 7B), two metrics for leakage in unparcellated/parcellated data respectively. However, choosing MNE/wMNE comes at the cost of localization errors (i.e., non-zero LE and fLE), a reduced robustness to sensor noise compared to eLORETA, and reduced ability to explain sensor-level data. These results are in line with those of dipole simulations, in which eLORETA largely outperformed MNE with the exception of a measure of leakage (‘focal width’) (Halder et al., 2019). It should additionally be noted that MNE/wMNE performed poorly on parcellated data (i.e., low 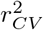, Figure 7). Therefore, although wMNE might be a useful tool if high spatial resolution is a priority, we recommend in this case that the use of eLORETA with alternative methods to reduce the effects of leakage such as orthogonalization (Colclough et al., 2015) is a more desirable approach.

Furthermore, source reconstructions from sLORETA (Pascual-Marqui, 2002) and the LCMV/wLCMV beamformers (Van Veen et al., 1997; Hillebrand et al., 2012) were less altered by the addition of Gaussian noise than eLORETA (Figure 5A), and explained less sensor-level data in low SNR (i.e., 100+ Hz) or zero SNR (i.e., pure white noise, Figure 5C) conditions. Therefore, sLORETA and beamformers may be appropriate if signal contamination from measurement noise or muscle artifacts is of concern. These results are in line with theory: sLORETA is unbiased in the presence of arbitrarily structured measurement or biological noise (Pascual-Marqui, 2007), and LCMV limits the influence of noise by minimizing variance from sources outside of the dipole of interest (Van Veen et al., 1997). Point dipole simulations have also verified that beamformers are insensitive to changes in SNR whilst minimum norm type estimates are not (Hincapié et al., 2017). However, the use of sLORETA/wLCMV comes at the cost of a significant drop in resolution compared to eLORETA (Figure 3E). This is particularly of concern in parcellated data (Figure 7B), as on average neighbouring ROIs had an instantaneous correlation of 78% (compared to eLORETA’s 46%), suggesting that sLORETA/wLCMV solutions lose finer functional details in anatomically adjacent regions, and hence these algorithms should be used with a more coarse grained atlas than the one in the current study. Whilst applying multivariate orthogonalization (Colclough et al., 2015) may address this issue, with such a large correlation between neighbouring regions there is a risk that much information will be lost when doing so.

Crucially, our findings highlighted that (*unweighted*) MNE (Hämäläinen and Ilmoniemi, 1994) and (*unweighted*) LCMV (Van Veen et al., 1997) are not appropriate for distributed, whole brain cortical reconstruction of restingstate MEG. MNE invariably explained 100% of the input data, but was unable to predict data from a left-out sensor in the cross validation procedure, suggesting an overfitting to the data that is not necessarily representative of true underlying source dynamics. LCMV exhibited high LE in line with studies using simulated dipoles (Pascual-Marqui et al., 2018; Halder et al., 2019), and was consistently outperformed by its weighted counterpart. Therefore for resting-state MEG source reconstruction, wLCMV is preferred over LCMV. Hillebrand et al. (2012) suggested that wLCMV should be used for resting-state data over LCMV due to the depth bias of the latter resulting in a non-uniform projection of sensor noise, potentially explaining our findings. Recently, an empirical Bayesian version of the LCMV beamformer (Belardinelli et al., 2012) has been shown to explain approximately 80% of crossvalidated variance in resting-state data (Little et al., 2018), which is still lower than that of eLORETA, but notably higher than either the classical LCMV or wLCMV, and hence could be a promising choice of beamformer-based solution.

One interesting finding of the current study was that, analytically, sLORETA and wLCMV produce non-identical, but similar solutions (Appendix C). Numerically, the differences in the performance of the two algorithms, whilst sometimes significant, were small in magnitude: wLCMV explained slightly more of the data (< 0. 1% difference in 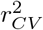), whilst sLORETA had marginally lower SECT (of the order 10^−3^mm difference). Therefore, under the conditions presented here, the two algorithms have negligible difference in practice. However, there may be cases where one solution is preferred over the other. One such case is when only a portion of the source space is to be reconstructed, for example placing a small number of ‘virtual electrodes’ at regions of interest (Engels et al., 2016; Hillebrand et al., 2016). wLCMV would be preferred in this case, as the beamformer solution at one dipole is not dependent on the rest of the source space (Van Veen et al., 1997). Conversely, when source reconstructing to a very high spatial resolution, sLORETA may be preferred over wLCMV due to two reasons. First, sLORETA had marginally better resolution statistics. Second, we proved analytically that, unlike sLORETA (Pascual-Marqui, 2002, 2007), wLCMV had small but non-zero LE, although such small LE from wLCMV could not be detected under the spatial grid used here. In fact, in the current study, wLCMV produced zero LE from empirical MEG data, consistent under eigendecomposition orientation constraints and after data parcellation. Future work could examine at what spatial resolution wLCMV produces non-zero LEs, and whether such high resolution is necessary or beneficial to inferences in the source space.

### 4.2. The reduced HCP-MMP atlas optimized for MEG

The original HCP-MMP atlas consists of 360 cortical ROIs delineated by their distinct structural, functional and connectivity profiles (Glasser et al., 2016), which offers good neuroanatomical precision essential for understanding macroscopic brain network dynamics. As such, the HCP-MMP atlas has been widely used for parcellation of resting-state fMRI data (Ito et al., 2017; Duboiş et al., 2018; Dermitaş et al., 2019; Preti and Van De Ville, 2019; Watanabe et al., 2019). Nevertheless, the fine spatial resolution of this atlas becomes a key limiting factor for apply it to MEG, because MEG data acquired in a typical scanner with 200-300 sensors would lead to rank deficiency if parcellated into 360 regions.

Here, we presented a reduction of the original HCP-MMP atlas with 250 cortical ROIs, in which deep regions with lower spatial resolution (chosen using a data-driven approach based upon the leadfield matrix, see Appendix B for details) were more coarse grained than in the original atlas. The target of reduction to 250 ROIs was determined because our MEG data consisted of 274 gradiometers, and after preprocessing (including noise projection and artifact rejection through ICA decomposition) had a rank of ≥ 250, although our method can be extended to reduce the original atlas further. Our approach posited that deeper regions of the brain or those with more radial orientation, which have low SNR (Cho et al., 2015) and low spatial resolution (Liu et al., 2002) for MEG, should contain fewer ROIs than superficial regions. This logic was supported by our spatial analysis of resolution metrics for all source reconstruction algorithms (Figure 4), where voxels with the low resolution (i.e. highest SECT) largely corresponded to those with low leadfield norm, quantified by a significant negative correlation between SECT and leadfield norm (Appendix B).

After determining the optimal number of ROIs per cluster, merging of the regions was done manually within each cluster, based on factors such as whether they were anatomical neighbours (required) and exhibited neuroanatomical and functional similarities (preferred) as reported by Glasser et al. (2016). Particularly, similarities in resting-state fMRI functional connectivity were prioritised, whilst properties such as myelination, cortical thickness, and activation during tasks were given low priority in the choice of ROIs to merge. Therefore, while the reduced atlas presented here is appropriate for parcellation of resting-state activity, one should use it with caution for task evoked MEG.

Alternative approaches exist for parcellating restingstate MEG data. First, one can opt for functionally derived ROIs, e.g. clustering voxels on the basis of functional connectivity profiles (for its application in fMRI data see Yeo et al., 2011). However, this approach is challenging for MEG data, due to the necessity of deciding between a large number of source reconstruction algorithms (section 2.1), functional connectivity metrics (Wendling et al., 2009; Dauwels et al., 2010; Wang et al., 2014) and frequency bands (Buzsáki, 2006), each combination of which likely results in a different functional connectivity profile (Hassan et al., 2014, 2017).

Second, one may use atlases for cortical parcellation with various levels of granularity, such as the Destrieux (74 cortical ROIs; Destrieux et al., 2010), AAL (90 cortical ROIs; Tzourio-Mazoyer et al., 2002), Desikan-Killiany (168 cortical ROIs; Desikan et al., 2006) and Brainnetome (210 cortical ROIs; Fan et al., 2016) atlases. The reduced HCP-MMP atlas presented here is advantageous for MEG research in two aspects: (1) it maintains all macroscopic-level clusters allowing for multi-modal comparisons with fMRI data; and (2) it maintains the fine grained resolution of the original HCP-MMP atlas for regions with high SNR in MEG data, whilst reducing the resolution only in deeper or more radial regions with low SNR in MEG data (Cho et al., 2015; Liu et al., 2002). Therefore, in superficial cortical regions where the MEG estimate is more reliable, the reduced HCP-MMP atlas has higher resolution than the above coarse grained atlases.

Finally, if MEG resting-state functional connectivity is the aim of the study, an additional alternative is to use the full HCP-MMP atlas with connectivity metrics insensitive to leakage, such as imaginary part of coherence (Nolte et al., 2004) or (weighted) phase lag index (Stam et al., 2007; Vinck et al., 2011). For standard connectivity metrics, orthogonalization between pairs of ROIs (Hipp et al., 2012) could be used, which does not require full rank data. However, multivariate leakage correction is preferred due to higher reliability of connectivity estimates than pairwise leakage correction (Colclough et al., 2015, 2016), and unlike pairwise correction does not exhibit spurious ‘ghost’ connections (Palva et al., 2018). Therefore, we suggest that the combination of our reduced HCP-MMP atlas and multivariate leakage correction is a more robust method to estimate resting-state MEG functional connectomes, as the reduced atlas also takes into account different signal-to-noise ratios in MEG data between brain regions. It is worth noting that multivariate leakage correction risks removing zero lag connectivity that is not an artifact of leakage. Recently, Farahibozorg et al. (2018) presented a split-and-merge algorithm to downsample anatomical atlases which alleviates the problem of leakage by merging ROIs based on CTFs derived from the resolution matrix (as opposed to norms of the leadfield presented here). This is potentially an advantage of the adaptive parcellations of Farahibozorg et al. (2018) over the method presented here. Nevertheless, since the CTF depends on the inverse solution, the resulting parcellation from that procedure is likely to also depend on the choice of source reconstruction algorithm. Since the current study suggested the use of eLORETA for parcellation based analyses, it would be of interest for future work to compare the reduced atlas presented here and an atlas derived following Farahibozorg et al. (2018) using the HCP-MMP atlas as a start point and eLORETA to construct CTFs.

### 4.3. Methodological considerations

A crucial choice that needed to be made for this study was the selection of source reconstruction algorithms to be compared. The list of algorithms studied here is by no means exhaustive, but instead focused on algorithms that may be useful for resting-state MEG data. Algorithms aimed towards localizing the spatial origin of a small number of active sources (i.e. *source localization*) were excluded, because this is likely to be an unrealistic assumption for resting-state. Examples of such source localization algorithms include dipole fitting (Scherg, 1990), multiple signal classification (Mosher and Leahy, 1998), and minimum norm estimation using multiple sparse priors (MSP, Friston et al., 2008; Little et al., 2018). Furthermore, we did not include algorithms that may perform well for source reconstructing resting-state data but are not compatible with our variance explained or resolution analyses, because comparison with the linear inverse algorithms presented here was not possible. These include algorithms which directly estimate resting-state source space functional networks from the sensor space data without first inverting the data, e.g. partial canonical coherence (Schoffelen and Gross, 2009; Popov et al., 2018), cortical partial coherence (Barzegaran and Knyazeva, 2017), and estimation of MVAR coefficients (Gómez-Herrero et al., 2008), as well as linear algorithms estimated in the frequency domain, e.g. dynamic imaging of coherent sources (Gross et al., 2001) and frequency domain minimum norm estimates (Yuan et al., 2008).

Of the source reconstruction algorithms most commonly used for resting-state analysis, the majority are covered in the current study. Notable exceptions include dynamic statistical parameter mapping (dSPM) (Dale et al., 2000), LORETA (Pascual Marqui et al., 1994), and the empirical Bayesian beamformer (EBB) (Belardinelli et al., 2012). Comparisons between dSPM and sLORETA have been studied using resolution metrics (Hauk et al., 2011; Hedrich et al., 2017; Hauk et al., 2019) and simulated dipoles (Pascual-Marqui et al., 2018), generally finding that sLORETA outperforms dSPM in terms of localization error, leakage, and false positive activity. Similarly, in a variance explained analysis of resting-state MEG, LORETA performed similarly to the MNE solution (Little et al., 2018). The EBB solution is derived from the LCMV beamformer, but is placed in a Bayesian framework in which hyperparameters of the inversion are optimized to increase model fit to the data (Wipf and Nagarajan, 2009; Belardinelli et al., 2012), meaning the variance explained analysis of EBB is not directly comparable with the other algorithms presented here. For these reasons, and to reduce the number of comparisons, dSPM/LORETA/EBB were excluded due to the inclusion of sLORETA/MNE/LCMV (respectively) in this study.

Another crucial methodological decision was choice of methods used to compare different algorithms. Previous studies have compared algorithms for *source localization* - identifying the origin of a small number of sources (Bai et al., 2007; Hassan et al., 2014; Bradley et al., 2016; Finger et al., 2016; Barzegaran and Knyazeva, 2017; Hassan et al., 2017; Hincapié et al., 2017; Bonaiuto et al., 2018; Pascual-Marqui et al., 2018; Seeland et al., 2018; Anzolin et al., 2019; Halder et al., 2019), such as known networks during task or simulated dipoles. These methods are not directly generalizable to resting-state data, where activity is not a point source but is distributed widely across the cortex. Instead, for resting-state data, one can ask to what extent true activity at a certain source location is mislocalized to another point on the cortex. Here, we approached the question of source (mis-)localization in resting-state data by considering resolution matrices (Hauk et al., 2011; Hedrich et al., 2017; Hauk et al., 2019). These matrices are the theoretical linear transformation between the true source activity and the estimated activity, and because of the linearity of this transformation results do not depend on number of active sources. For this reason, resolution metrics are equally as applicable to resting-state data as they are to evoked data.

There exists a large number of resolution metrics each of which can be applied to either the PSF or CTF (Hauk et al., 2019). The current study focused on two metrics (i.e., LE and SECT) to address crucial questions of interest. Firstly, given true (possibly distributed) activity on the cortex, how accurate is the placement of this activity in the source estimate? This question can be addressed quantitatively by ‘localization accuracy’ metrics applied to the PSF (Hauk et al., 2019), for which dipole localization error (LE) was measured in this study (Hauk et al., 2011; Hedrich et al., 2017; Hauk et al., 2019). Care must be taken when interpreting the LE in the resting-state paradigm. We are not attempting to quantify the difference in peak activation of the estimate given a single ‘true’ active dipole as in simulation or theoretical studies (Pascual-Marqui, 2007; Barzegaran and Knyazeva, 2017; Pascual-Marqui et al., 2018; Halder et al., 2019), because of the expected cross talk from distributed sources in resting-state data. As a result, ‘true’ activity at a certain location does not necessarily imply a peak of estimated activity at that location - for example, localization statistics based on peak activation for sLORETA are accurate for single dipole localization, but are imperfect for localizing multiple dipoles (Bradley et al., 2016). The LE statistic used in this study is therefore more properly interpreted as follows: given true activity at a dipole, that activity will most strongly influence dipoles approximately LE mm from the true source on average, regardless of activity at other locations.

The second key question we chose to address with resolution metrics was: given an estimate of activity at a given location, how strongly is this estimate influenced by leakage from other locations? This issue is important for resting-state functional connectivity analysis (Colclough et al., 2016), where spurious functional connections may arise due to leakage (Colclough et al., 2015). The latter can be addressed by ‘spatial extent’ metrics applied to the CTF (Hauk et al., 2019). Here, we chose a metric which has been called ‘spatial dispersion’ (SD) in past studies (Hauk et al., 2011; Hedrich et al., 2017; Hauk et al., 2019). When applied to the PSF this metric describes how activity at a given location ‘disperses’ due to leakage (hence the name SD), but when applied to the CTF this metric describes the spatial extent to which other locations influence the seed vertex through leakage. Hence, the name ‘spatial extent of cross talk’ (SECT) was used here to avoid confusion in interpretation, since in this study we do not quantify how activity at the seed disperses.

The aim of the resolution analysis is to compare the estimated source activity with the true source activity, based on the linear transformation that theoretically relates them. A complementary method of comparing the solutions would be to test the variance of source data explained by the estimate. This is usually not possible for empirical MEG data since the true source dynamics is unknown in the absence of simultaneous intracranial recordings. However, because the forward mapping has a unique solution for given source dynamics, (cross-validated) variance explained 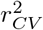 between the forward mapped source solution and the measured MEG data can be viewed as a proxy for accuracy of the source space solution. Indeed, Bonaiuto et al. (2018) found high correlations (0.98-1) between cross-validated sensor space errors and source space free energy for simulated dipoles. Sensor space variance explained is particularly useful for quantifying the quality of source reconstruction of resting-state data, since it makes no assumptions on the number or locations of active sources (i.e. can be applied to distributed cortical activity), is a whole brain measure as opposed to studying an individual ROI (Bonaiuto et al., 2018), and considers not only localization but also accuracy of the reconstructed time courses.

Finally, some settings in the source reconstruction pipeline may affect our results. Examples include the method used (including level of detail) for constructing the forward model (Hallez et al., 2007), the use of template models instead of subject specific models (Fuchs et al., 2002; Henson et al., 2009), and the density of the source space (Henson et al., 2009). In particular, the resolution metrics are likely to be dependent on these factors, due to the dependence of the resolution matrix on the leadfield matrix. Errors in the forward model are additionally known to affect the cross-validated variance explained (Little et al., 2018). By extension, since MEG and EEG give complementary information (Ding and Yuan, 2013) and the forward model is constructed differently between these modalities (Mosher et al., 1999), it should not be taken for granted that the results presented here apply to EEG resting-state data. Hence, the generalizability of our results to different acquisition modalities and robustness to differences in the forward model is an open question. Importantly, in this study the methodology was consistent across participants, and all statistics were performed within participants. Therefore, our comparisons between algorithms are unlikely to be biased or influenced by such confounding factors.

### 4.4. Conclusions

In conclusion, at both the voxel level and the ROI level, we recommend eLORETA (Pascual-Marqui, 2007, 2009) in combination with our new MEG-optimized reduction of the high resolution HCP-MMP atlas (Glasser et al., 2016) as an appropriate methodology for cortical source reconstruction of resting-state MEG. These conclusions are supported by quantitative evaluation demonstrating eLORETA has high capacity to explain the data even following parcellation, zero error in peak localization of cortical activity, and low leakage between ROIs compared to other algorithms.

## Acknowledgements

This study was supported by European Research Council (716321). AO was supported by a PhD studentship from Turkish Ministry of National Education. MJS was supported by a PhD studentship from Cardiff University School of Psychology. We thank Krish Singh and Alexander Shaw for helpful comments.

## Appendix

### A. Mathematical formulation of source reconstruction algorithms

In this appendix, we provide the mathematical framework for the source reconstruction algorithms used in the manuscript. All algorithms involve estimating the spatial filter **K** in Equation 2, which is an inverse of the leadfield matrix **Φ** in the forward model (Equation 1).

#### A.1. Beamformers

Beamformers calculate the filter for each dipole individually. Let us denote the column of the leadfield matrix corresponding to a dipole *i* as 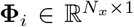. Then we define the beamformer weights for this dipole as 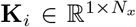, which can be used to construct a spatial filter **K** with rows **K**_*i*_.

##### A.1.1. LCMV beamformer

The linearly constrained minimum-variance (LCMV) beamformer (Van Veen et al., 1997) makes two constraints. Firstly, to ensure that **K** is the inverse of **Φ** and has unit output, we require **K**_*i*_**Φ**_*i*_ = 1. Secondly, to ensure zero response at other locations in the source space, we wish to attain **K**_*i*_**Φ**_*j*_ = 0 for *i* ≠ *j*. In practice this second constraint cannot be met exactly, but instead an optimal spatial filter can be designed which minimizes the variance of the output of the filter whilst still satisfying the first constraint. Mathematically, this problem is to minimize tr 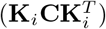, where **C** is the data covariance, subject to **K**_*i*_**Φ**_*i*_ = 1, and has the solution

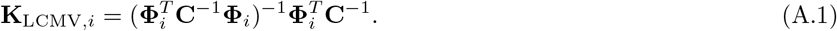

##### A.1.2. wLCMV beamformer

The LCMV filter is known to mislocalize superficial sources to deep locations due to non-uniform projections of sensor noise (Hillebrand et al., 2012). To account for this, the beamformer weights can be normalized to unit vector norm (Hillebrand et al., 2012). Here, we call this solution the *weighted LCMV* (wLCMV), given by

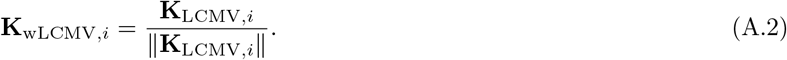

#### A.2. Minimum norm type estimates

Whilst beamformers consider each source point individually, regularized minimum norm type solutions reconstruct all sources within the source space simultaneously. An unregularized minimum norm estimate aims to minimize ||x — **Φ**ŝ||^2^, which has the Moore-Penrose psuedo-inverse **K** = **Φ**^*T*^(**ΦΦ**^*T*^) as a solution. However, since the problem is typically ill-posed, with *N_s_* ≫ *N_x_*, often this matrix **K** is singular. Regularization is often used to alleviate this issue, instead minimizing ||x – **Φ**ŝ||^2^ + λ**x**^*T*^**W**x. Here, λ = 0.05 is a regularization parameter which controls the extent to which the solution is regularized, and **W** is a weight matrix corresponding to a prior estimate of source variance-covariance (Dale et al., 2000; Hassan et al., 2014). The solution to this minimization is

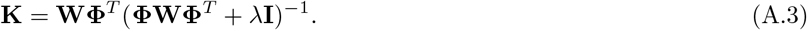

The MNE, wMNE, and eLORETA solutions all take this form, differing only in the prior estimate of source covariance **W**.

##### A.2.1. MNE

The solution often known simply as the minimum norm estimate (MNE; Hämäläinen and Ilmoniemi (1994)) makes a uniform estimate of source covariance, and is given by

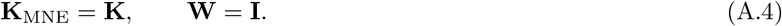

##### A.2.2. wMNE

The MNE solution suffers from depth bias, so the *weighted minimum norm estimate* sets the diagonals of **W** inversely proportional to the norm of the leadfield (Fuchs et al., 1999; Lin et al., 2006), i.e.

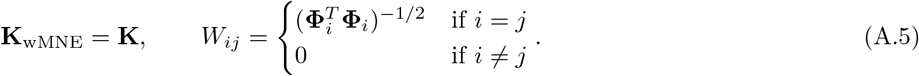

This solution essentially assumes as a prior that sources which only weakly influence the M/EEG must have a higher variance to be picked up by the sensors.

##### A.2.3. eLORETA

The *exact low resolution electromagnetic tomography* (eLORETA) solution (Pascual-Marqui, 2007, 2009) optimizes the weights such that not only is depth bias accounted for, the solution attains theoretically exact localization (Pascual-Marqui, 2007). The eLORETA solution is given by

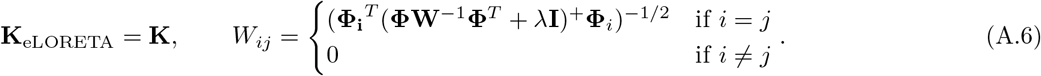

An iterative algorithm, described by Pascual-Marqui (2007), can be used to find these weights numerically.

##### A.2.4. sLORETA

Finally, an alternative approach to account for depth bias in the source localization is to estimate standardized distributions of current density, normalized by expected variance of each source. *Standardized low resolution electromagnetic tomography* (sLORETA; Pascual-Marqui (2002)) first calculates ŝ = **K**_MNE_*x* and then normalizes each time point to standardized variance. The sLORETA solution is therefore

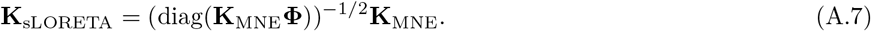

Whilst this standardization reduces the depth bias, the resulting solution can no longer be interpreted as a measure of the intensity of current density, and is more appropriately interpreted as probability of source activation.

### B. The reduced HCP-MMP atlas

#### B.1. Algorithm to identify the optimum number of ROIs per cluster

Below, we show a MATLAB code demonstrating the algorithm to identify the optimum number of ROIs per cluster. The strength of each of the 22 clusters Ω (i.e. ||**Φ**_Ω_|| in Equation 10 and normphiW in the code below; a 22 × 1 vector), the target number of regions (`targetN` in the code below; a scalar, here 125), and the number of ROIs per cluster in the original atlas (`n_orig`; a 22 × 1 vector) should be taken as input.

~~~
function n_new = determine_nRŨIs_per_cluster(normphiW,n_orig,targetN)
% Normalize the strengths of the regions to unit sum
normphiW = normphiW/sum(normphiW) ;
% Ideally, the number of RŨIs per region is proportional to the strength
n_new = round(targetN*normphiW) ;
% We don’t wish to split regions, only merge. Therefore we don’t want any
% regions to have a larger value in n_new than in n_orig
i_more = find(n_new > n_orig) ; % List of regions with larger value in % n_new than in n_orig
while ~isempty(i_more) % While there are regions with too many ROIs
        % Reset those regions to have the original number of ROIs
        n_new(i_more) = n_orig(i_more) ;
        % Recalculate target number of RŨIs, ignoring those we are fixing
        % to the original value
        targetN = targetN − sum(n_orig(i_more)) ;
        % Re-normalize strengths of RŨIs to unity, ignoring those we are
        % fixing to the original value
        normphiW(i_more) = nan ;
        normphiW = normphiW/nansum(normphiW) ;
        % Recalculate number of RŨIs in regions we are not setting to the
        % original value
        n_new = round(targetN*normphiW) ;
        n_new(i_more) = n_orig ;
        % Now we have a new n_new, once again calculate list of regions
        % with larger n_new than n_orig
        i_more = find(n_new./n_orig > 1) ;
end
~~~

#### B.2. Reducing the atlas

The algorithm above gives a target number of ROIs per cluster. To attain this target, we then examined each cluster and manually merged ROIs until the target number was achieved. To do so, the following criteria were used (in order of priority):

1. Merged ROIs *must* be in the same cluster.
2. Merged ROIs *must* be anatomical neighbours.
3. The resulting strengths of the merged ROIs should be approximately uniform. For example, consider a cluster with three ROIs, each of which neighbour both other ROIs (i.e. criteria 1 and 2 are satisfied). The ROIs have strengths 0.8, 0.4, and 0.3, and the target number of ROIs is two. We prefer to merge the ROIs with strengh 0.4 and 0.3 (resulting in ROIs with strengths 0.8 and 0.7) than, for example, the ROIs with strength 0.8 and 0.4 (resulting in ROIs with strengths 1.2 and 0.3). This result is justified, since we found from our resolution analysis that all algorithms demonstrated significant negative correlations between leakage (SECT) and the leadfield norm (LCMV/wLCMV/sLORETA *r* = –0.29; MNE *r* = –0.66, wMNE *r* = −0.69, eLORETA *r* = –0.68), suggesting that weaker sources have lower resolution, and therefore weaker ROIs should be merged to form larger but stronger ROIs.
4. Merged ROIs should display neuroanatomical and functional similarities, particularly having a low gradient in resting-state functional connectivity on the boundaries between the ROIs. These decisions were based on the detailed neuroanatomical descriptions given in Supplementary Material 3 of Glasser et al. (2016), in which a wide range of structural and functional imaging modalities were described for each cluster.

Supplementary Text S1 contains a list of all regions in the reduced atlas, and the choice of which ROIs were merged.

### C. Theoretical results relating sLORETA and wLCMV

In section 3.1, we found that the sLORETA and wLCMV results were similar, varying only by 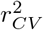 values of the order 10^−4^ to 10^−3^. In this section, we explore the theoretical relationship between the two methods. Interestingly, our numerical results suggested that wLCMV has zero localization error. Here, we explore this result further and demonstrate that, theoretically, wLCMV has non-zero localization error, and that our numerical results are potentially due to an insufficently coarse grid to identify these errors.

#### C.1. sLORETA and wLCMV have similar, but non-identical solutions

We begin with the sLORETA solution, given in Equation A.7. For dipole *i*, this can be written as

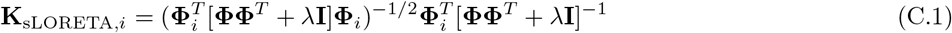

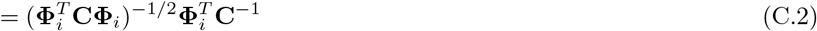

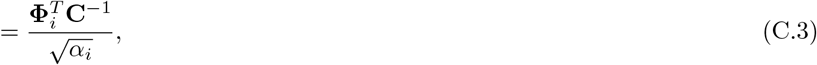

where

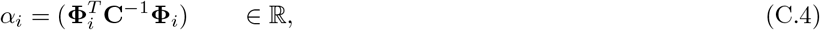

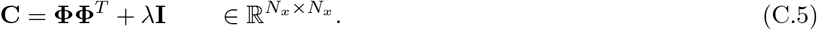

Here, 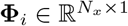 is the *i*th column of the leadfield.

Now consider the wLCMV solution, given by Equation A.1-A.2. We can write Equation A.1 as

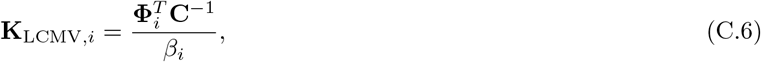

where here 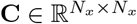 is the data covariance matrix and

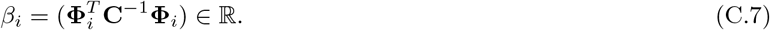

Therefore we can write Equation A.2 as

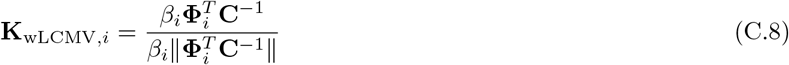

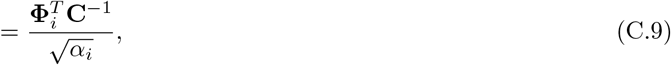

where

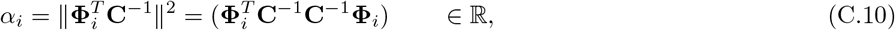

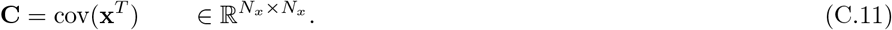

Here 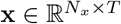 is the sensor space data.

By comparing Equation C.3 to Equation C.9, we can see that both solutions can be written in an identical form, where the differences lie in the scaling factor α_*i*_ and the matrix **C**, given by Equation C.4-C.5 for sLORETA and Equation C.10-C.11 for wLCMV. Namely, *α_i_* contains an addition factor of (**C**^−1^)^T^ in the wLCMV solution vs the sLORETA solution. In both cases the matrix **C** is a covariance matrix in sensor space. wLCMV uses the empirical M/EEG covariance matrix for **C**. sLORETA uses for **C** the theoretical M/EEG covariance matrix under the assumption of identity source covariance (including, if present, biological noise) ss^*T*^ = **I**, which maps to sensor space as **Φ_ss_**^*T*^**Φ**^*T*^ = **ΦΦ**^*T*^, and IID sensor noise with covariance λ**I**.

#### C.2. wLCMV has a theoretically non-zero localization error

Our derivation of exact localization is based around the resolution matrix defined in Equation 6 and follows that given to demonstrate exact localization of sLORETA and eLORETA by Pascual-Marqui (2007). Here, the equations are given in the case of known dipole orientations and in the absence of measurement noise, but the more general case of unconstrained dipoles and measurement/biological noise are given in Pascual-Marqui (2007). Consider an active dipole *j*. Then from Equation 6, the estimate at dipole *i* due to this dipole *j* is given by

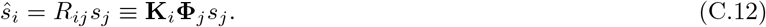

Here, 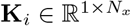 corresponds to the filter for the dipole *i*. Substituting the general form of the sLORETA and wLCMV solutions derived in Appendix C into this equation, we obtain

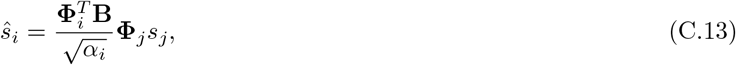

where for brevity we write **B** = **C**^−1^. The amplitude of the estimate at dipole *i*, *q_i_*, is therefore given by

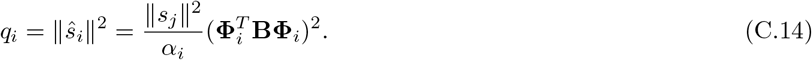

Our source localization methods attain zero localization error if the maximum of *q_i_* is at the original active dipole *j*. Therefore, following (Pascual-Marqui, 2007), we can write zero localization error is attained if

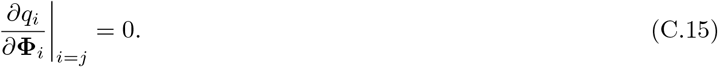

This derivative can be expressed as

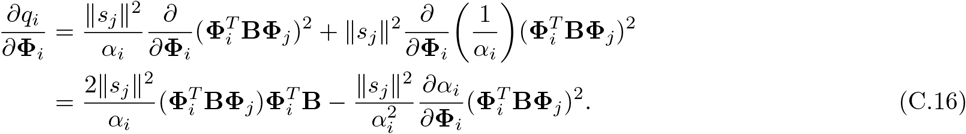

By writing Equation C.4 and C.10 in the same form, i.e.

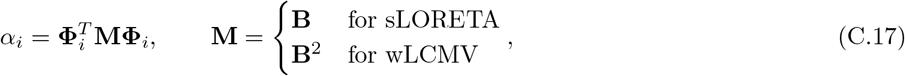

we can write

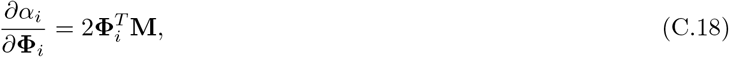

allowing us to explicitly write Equation C.16 as

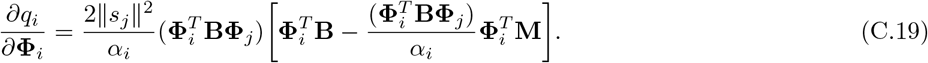

This derivative is zero at *i* = *j* iff

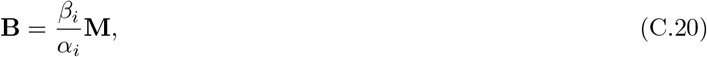

where *β_i_* is given in Equation C.7. For sLORETA, *α_i_* = *β_i_* (from Equation C.4 and Equation C.7), and **M** = **B** (Equation C.17), so it is clear that non-zero localization error is attained. For wLCMV, this is only true in the case where **B** = **B**^2^, which is true if **C**^−1^ = **I**. Therefore wLCMV only attains true zero localization error in the case where the data is IID.

To explain our numerical results that wLCMV solution showed zero localization error (Figure 3D), we studied Equation C.19 using our head model derived leadfields and empirical data covariance matrices. For each participant, 200 ‘active dipoles’ were randomly chosen without repetition (i.e. 200 unique values of *j* were chosen). For each value of *j*, we set *s_j_* = 1, and calculated the magnitude of *∂q_i_*/*∂***Φ**_i_ for *i* ≠ *j* using Equation C.19. To gain a normalized value to account for any global scaling, we additionally (for each value of j), drew 20 ‘incorrect’ dipoles with *i* ≠ *j*, recalculated the magnitude of *∂q_i_*/*∂*Φ_*i*_, and divided our raw ‘correct’ score by our average ‘incorrect’ score. Finally, we took the median over the 200 dipoles to obtain a single value per participant. We found *∂q_i_*/*∂***Φ**_*i*_ had a normalized magnitude of (2.19 ± 0.28) × 10^−14^. These results suggest that whilst *∂q_i_*/∂**Φ**_*i*_|*i*=*j* is non-zero for the wLCMV solution, it is close to zero (i.e. *i* = *j* is very close to a peak) and therefore the localization error is very small. Therefore, the zero localization error in numerical results of the wLCMV solution is likely due to the use of a discrete grid of spatial locations with spacing 4mm, making any localization errors undetectable. Such a grid spacing is common in the literature, so under these conditions wLCMV can be viewed in a practical sense as having zero localization error.

