## Supplementary Text and Figures for "Cortical source imaging of resting-state MEG with a high resolution atlas: An evaluation of methods"

### Supplementary Text S1. List of regions in the reduced HCP-MMP atlas

In the sections below, we describe which ROIs were merged and how this decision was made for each cluster. The order of the clusters described here matches those of Figure 6A and D from left to right, i.e. S1.4 Ventral stream visual cortex corresponds to purple in Figure 6A and D, since this is the fourth colour to appear from left to right. All neuroanatomical and functional descriptions used to guide our reduction, such as resting-state functional connectivity, myelination, cortical thickness, and activation during tasks, are taken from Supplementary Material 3 of Glasser et al. (2016).

#### *S1.1. Primary visual cortex (V1)*

In the original atlas, this first cluster contains only a single ROI, the primary visual cortex (V1). Regardless of the fact it only contains a single ROI, V1 is the strongest cluster. In the initial iteration of the algorithm, V1 was actually ideally predicted to contain four ROIs. However, since we are aiming to reduce the number of ROIs, we do not split ROIs. Therefore V1 was not altered in the reduced atlas.

#### *S1.2. Early visual cortex*

The early visual cortex contains the second, third, and fourth visual areas (V2, V3, V4). Like cluster 1, cluster 2 had very high strength, and in the initial iteration of the algorithm was predicted to have an optimum number of 7 ROIs. Accounting for the fact we do not split ROIs, the final iterate of the algorithm suggested three ROIs was optimum, and hence cluster 2 was unaltered in the reduced atlas.

#### *S1.3. Dorsal stream visual cortex*

In the original atlas, the dorsal visual stream contains six ROIs, including five visual areas (V3A, V3B, V6, V6A, V7) and an intraparietal sulcus area (IPS1). We found the strengths of each ROI to be in the range 0.2182% to 0.8123%. Our algorithm identified the optimum number of ROIs to be four. Across the dorsal stream visual cortex, there was relative homogeneity in resting-state functional connectivity, with primary differences between ROIs being in terms of myelination and activation during a battery of tasks. We therefore merged V3B (0.2182%) with its neighbour V7 (0.3485%; total strength 0.5556%), which have no resting-state FC gradient and differ from neighbouring areas V6A, IPS1, IP0 and V3CD in having more myelin. We additionally merged V6 (0.5222%) and V6A (0.2821%; total strength 0.8043%), which differed only in working memory and MOTOR AVG tasks. The resulting range of strengths was 0.4741% to 0.8123%. The resulting four regions were therefore V3A, V3B+V7, V6+V6A, IPS1.

#### *S1.4. Ventral stream visual cortex*

In the HCP-MMP atlas, the ventral visual stream contains seven ROIs including visual area 8 (V8), the ventral visual complex (VVC), the PIT complex (PIT), the fusiform face complex (FFC), and the ventro-medial visual areas 1-3 (VMV1, VMV2, and VMV3). We found the strengths of each ROI to be in the range 0.0843% to 0.5656%. Our algorithm identified the optimum number of ROIs to be three. Across the ventral stream visual cortex, there was relative homogeneity in resting-state functional connectivity, with primary differences between ROIs being in terms of functional activation during a battery of tasks. V8 (0.2741%) and VVC (0.3691%) were reported by Glasser et al. (2016) to be the heavily myelinated core of the ventral visual stream, differing only by a visuotopic boundary and cortical thickness, so we chose these two ROIs to merge (total strength 0.6432%). Glasser et al. (2016) additionally reported the only significant difference between PIT (0.3911%) and FFC (0.5656%) was for cortical thickness, so these two ROIs were merged (total strength 0.9567%). Finally, we merged the neighbouring ventro-medial visual areas VMV1 (0.1524%), VMV2 (0.0843%), and VMV3 (0.1160%; total strength (0.3527%). The resulting range of strengths was 0.3527% to 0.9567%. Whilst this is quite a broad range, we were limited by the anatomical organization of this cluster; VMV1-3 form a strip, with the only anatomical neighbours being VVC and V8 on the border of V3, whilst FFC and PIT lay on the opposite side of the VVC+V8 complex. This means achieving homogeneity in the strengths of the merged regions whilst achieving neuroanatomical and functional similarity was not possible. The resulting three regions were therefore V8+VVC, VMV1-3, PIT+FFC.

#### *S1.5. MT+ complex and neighbouring visual areas*

Cluster 5 contains nine ROIs including four in the MT+ complex (MT, MST, V4t, and FST), the adjacent area PH, three lateral occipital areas (LO1, LO2, and LO3), and their adjacent area V3CD. We found the strengths of each ROI to be in the range 0.2455% to 0.7601%. Our algorithm identified the optimum number of ROIs to be five. Homoegeneity of the MT+ complex in terms of resting-state functional connectivity has been reported; particularly the MT/MST boundary was reported to be less robust than most others reported by Glasser et al. (2016), differing primarily in terms of a single RANDOM-TOM task. The MST/MT regions additionally differed from the rest of the MT+ complex primary due to activation in the BODY-AVG task, we no difference in resting-state functional connectivity. We therefore merge MT (0.3534%), MST (0.2455%), and V4t (0.2767%; total strength (0.8765%). We also merged the three lateral occipital areas LO1 (0.2960%), LO2 (0.2828%), and LO3 (0.3805%; total strength 0.9593%), which differed in cortical thickness and a battery of tasks. The resulting range of strengths was (0.4868% to 0.9593%), and the resulting five regions were MT+MST+V4t, FST, PH, LO1-3, V3CD.

#### *S1.6. Somatosensory and motor cortex*

The somatosensory and motor cortex contains the cortical Brodmann areas 1, 2, 3a, 3b, and 4. The initial iterate of the algorithm predicted cluster 6 should contain 10 ROIs, so like clusters 1, 2, and 17, this cluster is actually over represented in the reduced parcellation. Accounting for the fact we do not split ROIs, the final iterate of the algorithm suggested five ROIs was optimum, and hence cluster 6 was unaltered in the reduced atlas.

#### *S1.7. Paracentral lobular and mid-cingulate cortex*

Cluster 7 contains eight ROIs including the dorsal and ventral cingulate motor areas 24d (24dd and 24dv), supplementary motor areas (6mp, 6ma, SCEF), and subdivisions of area 5 (5m, 5L, 5mv). The strengths of these ROIs were in the range 0.2163% to 0.9348%. Our algorithm identified the optimum number of ROIs to be 7. Here, we considered the distributions of strengths of ROIs. The supplementary motor areas had strengths in the range 0.7553-0.9348%, whilst area 5 had strengths in the range 0.3799-0.6370, and area 24d had strenths 0.2163% (24dv) and 0.4519% (24dd) respectively. Therefore, whilst Glasser et al. (2016) reported a small difference in resting-state functional connectivity between 24dd and 24dv, this was also notably the weakest set of regions and hence in the interest of uniformity in ROI strengths we merged these two regions (total strength 0.6682%). The resulting range of strengths was 0.3799% to 0.9348%. The seven regions in the resulting merged atlas were therefore 24dd+24dv, 6mp, 6ma, SCEF, 5m, 5L, and 5mv.

#### *S1.8. Premotor cortex*

The premotor cortex contains seven ROIs including area 55b (55b), the superior premotor regions (6d, 6a) and frontal eye field (FEF), and the inferior premotor regions (6v, 6r) and premotor eye field (PEF). The strengths of these regions ranged from 0.3162% to 0.9661%. Our algorithm suggested this cluster should contain six ROIs. Whilst there were strong differences in resting-state functional connectivity, myelination, and task activation between 55b and PEF (Glasser et al., 2016), these two neighbouring regions were notably the weakest (strengths of 0.4480% and 0.3162% respectively). We therefore merged these regions (total strength 0.7642%) to maintain uniformity of ROIs, resulting in a range of strengths 0.5666% to 0.9661%. Therefore the resulting ROIs were 55b+PEF, 6d, 6a, FEF, 6v, and 6r.

#### *S1.9. Posterior opercular cortex*

The posterior opercular cortex contains six ROIs. These are area 43 (43), the frontal opercular area 1 (FOP1), opercular areas 1-4 (OP1, OP2-3, OP4), and area PFcm (PFcm). These regions range in strength from 0.1278% to 0.5843%. FOP1 (0.1799%) was one of the weakest areas and had a single anatomical neighbour 43 (0.3886%) with a small difference in functional connectivity and a large difference in activation during the motor task (Glasser et al., 2016), so we chose to merge these ROIs (total strength 0.5864%). We additionally merged areas OP1 (0.2752%), OP2-3 (0.1278%) and PFcm (0.3518%; total strength 0.7548%), Between these areas, the primary differences are in activation during tasks such as language and motor, cortical thickness, and a small gradient in functional connectivity. The resulting range of strengths was 0.5843% to 0.7548%.

#### *S1.10. Early auditory cortex*

Cluster 10 contains five ROIs including the primary auditory cortex (A1), the lateral, medial, and para- belts (LBelt, MBelt, PBelt), and the retro-insular cortex (RI). These regions range in strength from 0.1429% to 0.4705%. Our algorithm suggests this cluster should contain two ROIs. PBelt (0.4705%) and RI (0.1931%) has no gradient in resting-state functional connectivity along the border, differing primarily in in myelination (however, PBelt does demonstrate stronger functional connectivity with a thalamic seed), so these two ROIs were merged (total strength 0.6636%). Similarly, A1 (0.1429%) and MBelt (0.2102%) demonstrated no gradient in resting-state functional connectivity (although these regions differed in myelination, activation in the motor-cue task, and function connectivity with a thalamic seed), so these ROIs were also merged. Finally, LBelt (0.1835%) is anatomical neighbours with both PBelt+RI and A1+MBelt and has a moderate FC gradient along borders with both. Therefore we merge LBelt with A1+MBelt (total strength 0.5366%) to achieve uniformity in strengths. Our final ROIs were therefore A1+MBelt+LBelt and PBelt+RI.

*S1.11. Auditory association cortex*

Cluster 11 contains eight ROIs including auditory complexes 4 and 5 (A4, A5), the dorsal/ventral anterior/posterior superior temporal sulcus (STSda, STSdp, STSva, STSvp), the superior temporal gyrus region a (STGa), and temporal region A2 (TA2). These regions range in strength from 0.2579% to 0.7058%. Our algorithm predicted the target number of ROIs in this cluster was six. This region was reasonably homogeneous in resting-state functional connectivity, with the primary FC gradient falling on the dorsal/ventral border of the superior temporal sulcus. The two weakest ROIs were STGa (0.2579%) and TA2 (0.3189%), which are anatomical neighbours which lie on the planum polare and superior temporal gyrus anterior to Heschl's gyrus, with main differences in terms of activation during tasks (Glasser et al., 2016). We therefore merged these ROIs (total strength 0.5768%). The next weakest logical pair of ROIs was the STSda (0.4426%) and STSdp (0.3657%), which are anatomical neighbours on the dorsal superior temporal sulcus and differed only in terms of myelination and activation during tasks, with no differences in resting-state functional connectivity. Therefore these ROIs were also merged (total strength 0.8083%). The resulting strengths of ROIs were in the range 0.4634% to 0.8083%. The resulting six ROIs were A4, A5, STSda+STSdp, STSva, STSvp, STGa+TA2.

*S1.12. Insular and frontal opercular cortex*

In the original HCP-MMP atlas, the insular and frontal opercular cortex were very described in very high resolution, consisting of thirteen ROIs including area 52 (52), the parainsular cortex (PI), the insula granular (Ig), two posterior insular areas (PoI1 and PoI2), four frontal opercular areas (FOP2, FOP3, FOP4, and FOP5), the middle and anterior ventral insular areas (MI and AVI), the anterior agranular insular complex (AAIC), and the piriform cortex (Pir). However, due to the depth of this cluster, these ROIs had relatively low strength, ranging from 0.0724% to 0.2759%, with the exception of Pir which had a notably larger strength 0.6485%. Therefore our algorithm suggested this region should consist of only four ROIs.

Among the four frontal opercular areas, there is very little resting-state FC gradient, and differences are primarily in cortical thickness, gambling reward, math, and motor tasks (Glasser et al., 2016). Furthermore, in our analysis FOP2-3 were among the weakest ROIs (0.0889% and 0.0588% respectively), whilst FOP4-5 were among the strongest when Pir was excluded (0.2759% and 0.2483% respectively). We therefore merged these ROIs to form a single frontal opercular ROI (total strength 0.6719%).

Another group of ROIs that are largely homogeneous in terms of resting-state FC are Ig (strength 0.0660%), 52 (0.0724%), PoI1 (0.1220%), PoI2 (0.1255%), and PI (0.1854%). This group of ROIs has a strong gradient in resting-state FC on the PoI2 border with MI and AAIC and the Ig border with FOP2, but internally displays relatively little FC gradient. However, it is worth noting that 52 has a much stronger resting-state FC with a thalamic seed than the rest of this group. We therefore merge these ROIs (total strength 0.5713%).

The remaining ROIs which have not been merged at this stage are Pir, AAIC, AVI, MI. Since Pir was notably stronger than all other ROIs in this cluster, we did not merge it with any other ROIs. The final merged ROI therefore consisted of AAIC (0.0953%), AVI (0.1605%), and MI (0.1598%), which make up the middle and anterior region of the insular cortex (total strength 0.4156%). Unfortunately, in terms of resting-state FC, these regions are highly heterogeneous, both between and within the ROIs. However, due to the low resolution of MEG and the deep/weak nature of these regions, we propose that even a much higher resolution parcellation would be unlikely to identify these fine FC alterations as the insular cortex had some of the highest spatial extents of cross talk (Figure 4).

Our final four ROIs in cluster 12 were therefore Pir, FOP2+FOP3+FOP4+FOP5, Ig+52+PoI1 +PoI2+PI, and AAIC+AVI+MI, with strengths ranging from 0.4156% to 0.6719%.

*S1.13. Medial temporal cortex*

In the original HCP-MMP atlas, the medial temporal region contained seven regions; however here we considered only six of these regions, excluding the hippocampal grey matter from the analysis. These six ROIs were three perihippocampal areas (PHA1, PHA2, PHA3), the peri-entorhinal/ectorhinal complex (PeEc), the entorhinal cortex (EC), and the presubiculum (PreS). The strengths of these ROIs were in the range 0.0905% to 0.6193%, and our algorithm suggested there should be three ROIs within this cluster. There was a large resting-state FC gradient throughout this region, so our motivation for merging ROIs was predominantly motivated by aiming towards uniform strengths. The three perihippocampal areas were weakest (PHA1 0.1355%, PHA2 0.0905%, PHA3 0.1129%), so we first merged these into a single perihippocampal area (total strength 0.3489%). This was still the weakest ROI, and was neighbours with all other ROIs. Since PreS (0.5196%) and PeEc (0.6193%) were notably stronger than EC (0.3650%), we further merged PHA1-3 with EC (total strength 0.7139%). The resulting range of strengths was therefore 0.5196% to 0.7139%. The resulting three ROIs in the cluster were PreS, PeEc, and PHA1+PHA2+PHA3+EC.

*S1.14. Lateral temporal cortex*

The lateral temporal cortex contains a total of nine ROIs, including two (TGd and TGv) in the temporal polar cortex, three (TF, TE2a, and TE2p) in the inferior temporal sulcus and gyrus, and four (TE1a, TE1m, TE1p, and PHT) in the middle temporal gyrus. The algorithm suggested nine ROIs was optimum, and hence cluster 14 was unaltered in the reduced atlas.

*S1.15. Temporo-parieto-occipital junction*

Cluster 15 contains five ROIs, including three on the temporo-parieto-occipital junction (TPOJ1, TPOJ2, and TPOJ3), the Perisylvian language area (PSL), and the superior temporal visual area (STV). The optimum number of ROIs was five, so cluster 15 was unaltered in the reduced atlas.

*S1.16. Superior parietal cortex*

The superior parietal cortex contains ten ROIs, including medial and lateral area 7P (7Pm, 7PL), medial and lateral area 7A (7Am, 7AL), area 7PC (7PC), and five intraparietal/lateral intraparietal areas (LIPv, LIPd, VIP, MIP, AIP). These ROIs ranged in strengths from 0.2793% to 0.9229%. Our algorithm suggested the optimum number of ROIs was nine, so based primarily on strengths and FC gradients reported by Glasser et al. (2016), we chose to merge the lateral and medial areas 7P (7PL 0.3923%, 7Pm 0.3912%; total 0.7835%).

*S1.17. Inferior parietal cortex*

The inferior parietal cortex contains 10 ROIs, including a transitional area PGp, three areas on the intraparietal sulcus (IP0, IP1, and IP2), three areas corresponding to a subdivision of the classical PF (PF, PFt, and PFop), and three nodes of the task positive network (PFm, PGi, PGs). The initial iterate of the algorithm predicted cluster 17 should contain 13 ROIs, so like clusters 1, 2, and 6, this cluster is actually over represented in the reduced parcellation. Accounting for the fact we do not split ROIs, the final iterate of the algorithm suggested ten ROIs was optimum, and hence cluster 17 was unaltered in the reduced atlas.

*S1.18. Posterior cingulate cortex*

The posterior cingulate cortex contains 14 ROIs, which Glasser et al. (2016) described as functionally distinct, and grouped based upon geographical proximity rather than functional similarity. There are large resting-state functional connectivity gradients on the borders of many ROIs within this cluster, so uniform strengths were prioritised in merging ROIs for cluster 18. Strengths of the ROIs ranged from 0.0995% to 0.4940%, excluding the parietal occipital sulcus area 2 (POS2) which had a strength of 0.9410%. Our algorithm suggested the optimum number of ROIs was six.

The first merged ROI consisted of transitional areas between the early visual cortex and posterior cingulate association cortex found on the anterior bank of the parietal-occipital sulcus. These ROIs were the dorsal visual transitional (DVT; 0.4940%) and prostriate (ProS; 0.0998%) cortices (total strength 0.5938%), which differed primarily in myelination and activation during a number of tasks. Furthermore, Glasser et al. (2016) separated area 31 into three ROIs, namely anterior 31 (31a; 0.1261%) and dorsal/ventral posterior 31 (31pd/31pv; 0.1445%/0.1078% respectively). Borders between these regions were primarily due to activation in tasks, although modest gradients in myelination and resting-state functional connectivity were also reported. Therefore, for our second ROI we merged these three ROIs into a single area 31 (total strength 0.3784%).

Similarly to area 31, Glasser et al. (2016) subdivided area 23 into the dorsal and ventral 23ab (d23ab/v23ab; 0.1006%/0.0995%), 23c (0.2997%), and 23d (0.1579%). With the exception of 23c, these areas differed only modestly in resting-state functional connectivity. However, there was a strong gradient in resting-state functional connectivity along the border of 23c the rest of 23. We therefore merged d23ab, v23ab, and 23d, but did not include 23c in this ROI. Instead, 23c and the precuneus visual area (PCV; 0.4299%) were merged (total strength 0.7296%), based on the fact these ROIs did not differ in resting-state functional connectivity, but did have a strong gradient along their borders with the rest of cluster 18.

In order to maintain uniformity in terms of ROI strengths, the d23ab+v23ab+23d ROI was additionally merged with neighbouring area 7m (0.4078%), which had similar resting-state functional connectivity (total strength 0.7658%).

Three ROIs remain which have not been discussed in detail, namely parieto-occipital sulcus areas 1 (POS1; 0.2965%) and 2 (POS2; 0.9410%), and the restrosplenial complex (RSC; 0.4614%). Since POS2 had a notably larger strength than any other ROI, we did not merge this with any other ROIs. POS1 was merged with its neighbour RSC (total strength 0.7579%). The resulting range of strengths for cluster 18 were therefore 0.3784% to 0.9410%. The six resulting ROIs for this cluster were DVT+ProS, d23ab+v23ab+23d+7m, 31a+31pd+31pv, POS1+RSC, PCV+23c, POS2.

#### *S1.19. Anterior cingulate and medial prefrontal cortex*

Cluster 19 had 15 ROIs in the original HCP-MMP parcellation. Strengths of ROIs within this cluster ranged from 0.1028% to 0.8540%. Our algorithm suggested this cluster should contain eight ROIs.

Firstly, we considered the anterior cingulate cortex. Glasser et al. (2016) describes three classical bands of cingulate cortex corresponding to area 33, area 24, and area 32. Whilst these cingulate areas were reasonably homogeneous in resting-state functional connectivity, a strong gradient separated them from much of the dorsal medial prefrontal cortex. In area 33, (Glasser et al., 2016) described a single area 33pr (33pr; 0.4011%). Area 24 was separated into areas 24 and 24pr, which were subdivided into the anterior/posterior 24 (a24/p24; 0.4039%/0.3747%) and anterior/posterior 24pr (a24pr/p24pr; 0.1305%/0.1719%). Area 32 was divided into anterior/posterior 32pr (a32pr/p32pr; 0.1634%/0.2199%), d32 (0.2503%), p32 (0.1771%), and s32 (0.1028%). p32 and s32 were separated from the remaining areas by a band of medial prefrontal cortex 9m (discussed below). Following these descriptions, we merged the anterior cingulate cortex into the following ROIs: 33pr, a24pr+p24pr (total strength 0.3024%), a24+p24 (total strength 0.7786%), and a32pr+p32pr+d32 (total strength 0.6336%). Furthermore, p32+s32 were merged with the neighbouring subcallosal cingulate area 25 (0.3892%; total strength 0.6691%).

The medial prefrontal cortex consisted of an additional four ROIs in the original parcellation. The strongest ROIs in cluster 19 are the superior medial prefrontal areas 8BM (0.7088%) and 9m (0.8540%), which are separated from the cingulate regions and each other by a strong gradient in resting-state functional. On the medial surface of the orbital prefrontal cortex were two subdivisions of area 10, namely 10r (0.2384%) and 10v (0.4945%), which differ modestly in resting-state functional connectivity (Glasser et al., 2016). The two subdivisions of area 10 were merged (total strength 0.7779%). The resulting eight ROIs in cluster 19 therefore ranged in strength from 0.3024% to 0.8540%, and were 33pr, a24pr+p24pr, a24+p24, a32pr+p32pr+d32, p32+s32+25, 10r+10v, 8BM, and 9m.

#### *S1.20. Orbital and polar frontal cortex*

The orbital and prefrontal cortex consisted of 11 ROIs, including the orbitofrontal and posterior orbitofrontal complexes (OFC, pOFC), areas 13l and 11l, three subdivisions of area 47 (47s, 47m, a47r), and four subdivisions of area 10 (10pp, a10p, p10p, and 10d).

These ROIs ranged in strength from 0.1311% to 0.7791%. Our algorithm suggested seven ROIs for this region. Two of the weakest ROIs in this cluster were subdivisions of area 47, 47m (0.1311%) and 47s (0.2807%). Glasser et al. (2016) reported a small difference in functional connectivity between these regions, but the primary difference was in myelination. We therefore merged these regions (total strength 0.4118%). Interestingly, the remaining subdivision of 47, a47r (0.7791%) was the strongest ROI, so this was not merged with 47m+47s.

13l (0.2327%) was the next weakest ROI. We merged 13l with 11l (0.5175%; total strength 0.7502%), forming a complex which lies lateral to the orbitofrontal complexes (Glasser et al., 2016). We additionally merged the four subdivisions of area 10 into two pairs of subdivisions; the anterior/posterior area 10p (a10p/p10p; 0.4088%/0.5093%) formed one subdivision (total strength 0.9181%), whilst 10d (0.5302%) and 10pp (0.3362%) formed the other subdivision (total strength 0.8664%). The range of strengths in the resulting seven ROIs were 0.4118% to 0.9181%, which were OFC, pOFC, 13l+11l, 47s+47m, a47r, 10pp+10d, and a10p+p10p.

*S1.21. Inferior frontal cortex*

The inferior frontal cortex consists of eight ROIs including areas 44, 45, two subdivisions of 47 (47l, p47r), and four subdivisions of the inferior frontal sulcus (IFJp, IFJa, IFSp, and IFSa). Excluding the weak anterior/posterior IFJ regions (IFJa/IFJp; 0.2140%/0.1139%), these ROIs were largely heterogeneous in strength, ranging from 0.4280% to 0.6664%. Our algorithm suggested that this cluster should contain six ROIs, so IFJa and IFJp were merged with the neigbouring IFSp (0.4280%; total strength 0.7759%). The resulting range of strengths were therefore 0.4811% to 0.7759%. The six ROIs in cluster 21 were therefore 44, 45, 47l, p47r, IFJa+IFJp+IFSp, IFSa.

*S1.22. Dorsolateral prefrontal cortex*

The dorsolateral prefrontal cortex contains 13 ROIs, including the superiod frontal language are (SFL), two transitional areas between areas 6 and 8 (s6-8 and i6-8), a subdivision of area 8 into four areas (8BL, 8Ad, 8Av, and 8C), a subdivision of area 9 into two areas (9p and 9a), and four areas in the central portion of the dorsolateral prefrontal region (9-46d, 46, a9-46v, and p9-46v). The algorithm suggested 13 ROIs was optimum, and hence cluster 22 was unaltered in the reduced atlas.

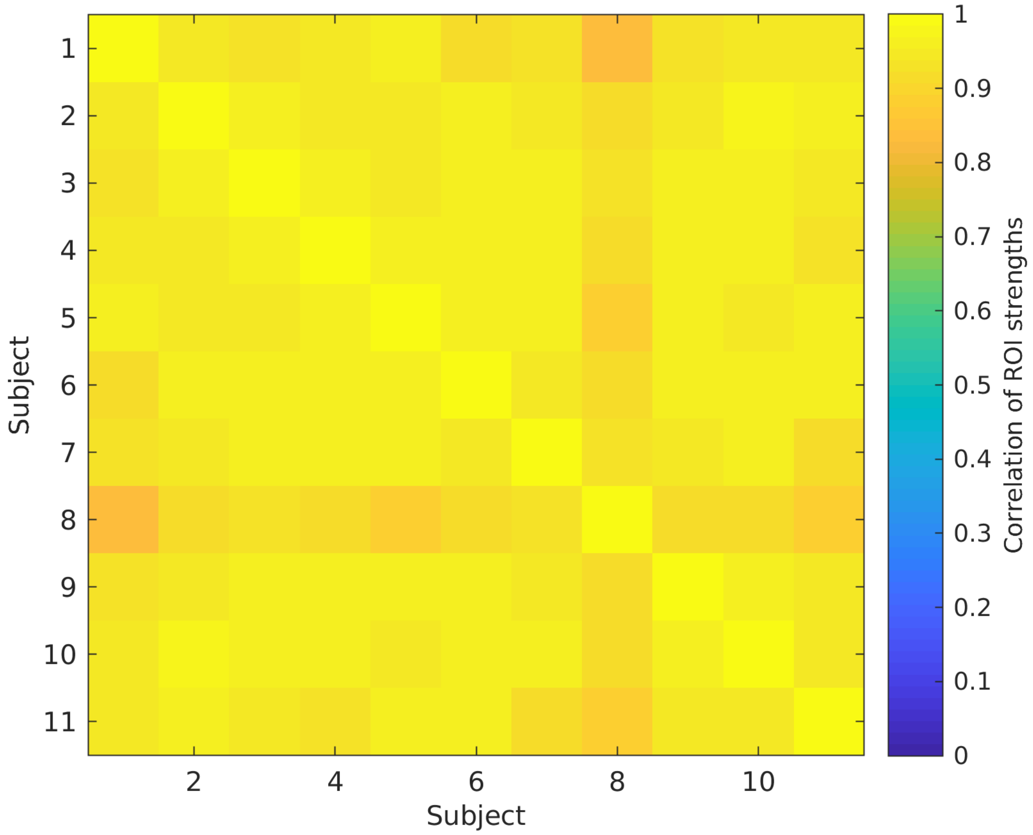


Figure S1: **ROI strengths are highly consistent across subjects**. The reduction of the HCP-MMP parcellation, described in section 2.3 and shown in Figure 6, made use of the average ROI strength for each ROI over 11 participants. To ensure these values were consistent across subjects, i.e. the same areas had large/small strength in all subjects, the distributions of ROIs strengths were correlated. All pairs of subjects had a correlation $> 0.83$, suggesting the spatial patterns of ROI strengths are highly consistent across subjects.

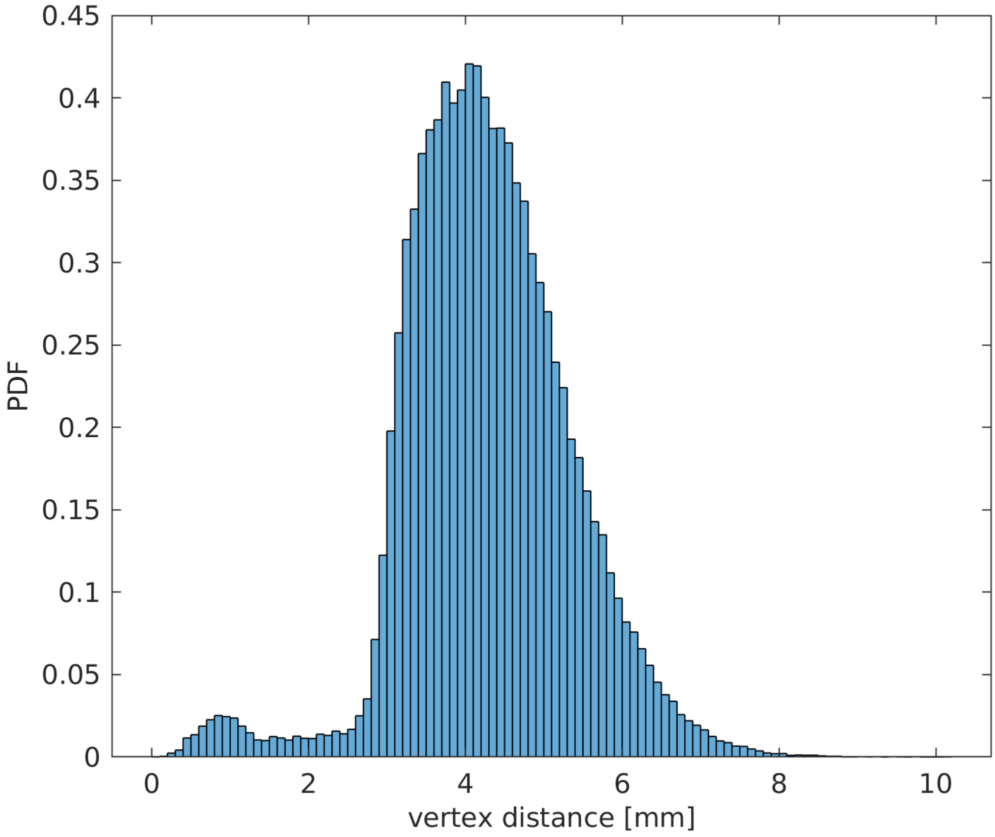


Figure S2: **Distribution of distances between neighbouring vertices in cortical mesh.** For each participant, 10,000 pairs of neighbouring vertices in the cortical mesh were randomly sampled without repetition and the distance between pairs calculated. Shown here is a histogram, normalized to a probability distribution function (PDF), of distances over the resulting 110,000 distances sampled.

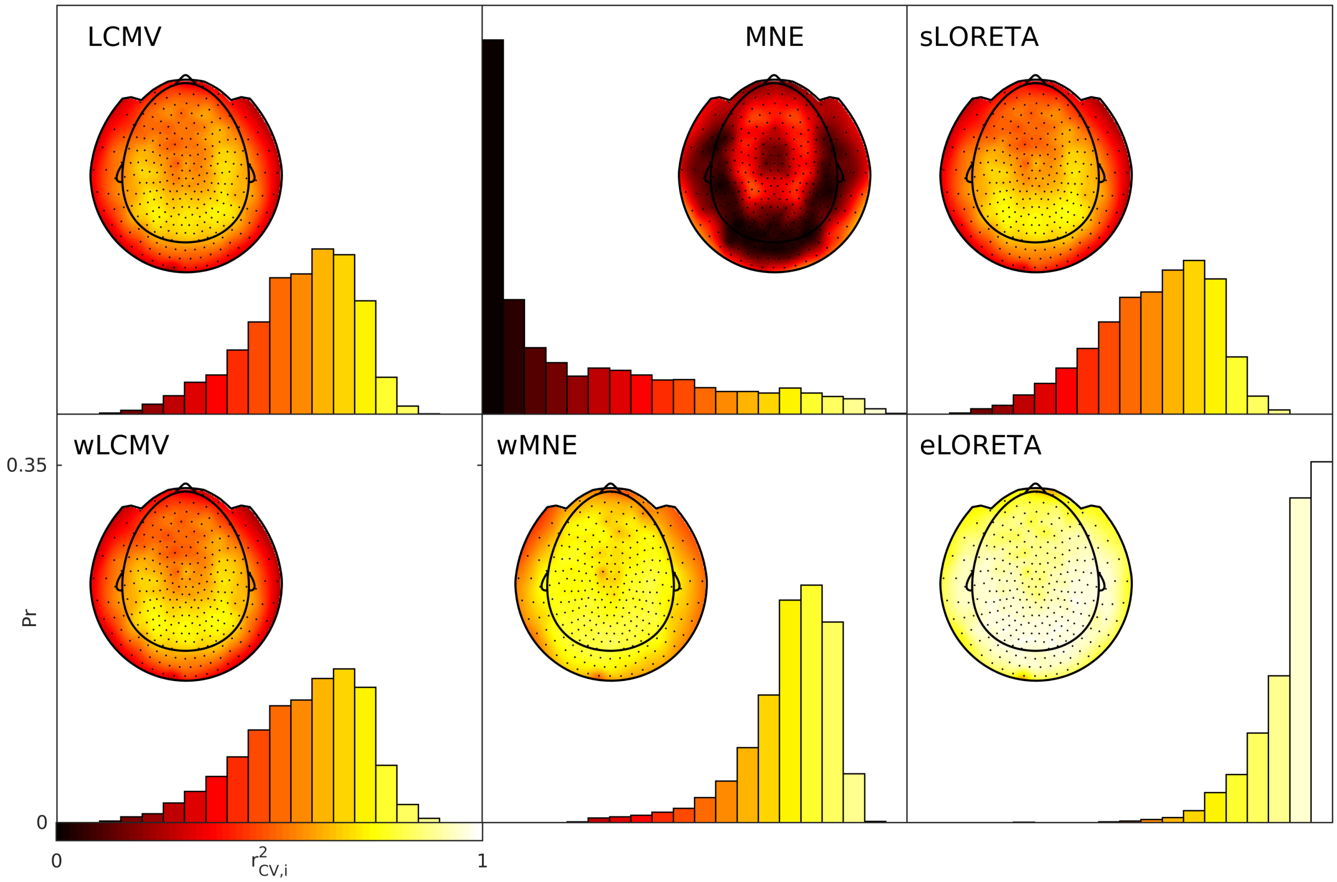


Figure S3: **Distributions of $r^2_{CV}$ values for all MEG sensors and participants**. The boxplots in main text Figure 3A show the averages of this data over MEG sensors for each participant. For each algorithm, horizontal and vertical axes and colour scale are identical and calibrated in the bottom left. Inlays show the spatial distributions of these values, averaged over participants. The colour scale is identical for the inlays and the histograms, shown in the bottom left.

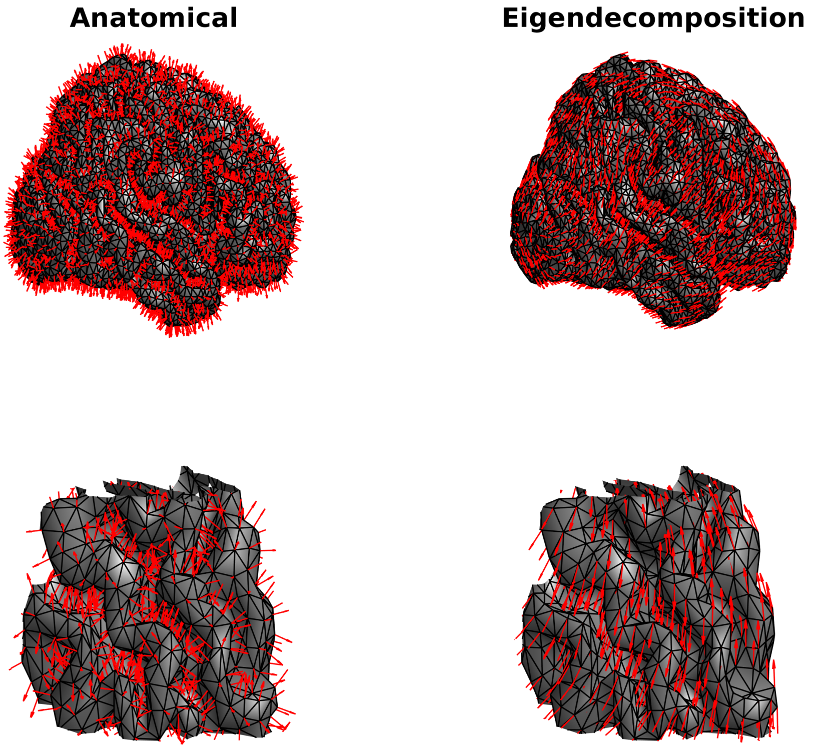


Figure S4: **Example of dipole orientations for a single participant using different orientation constraints.**

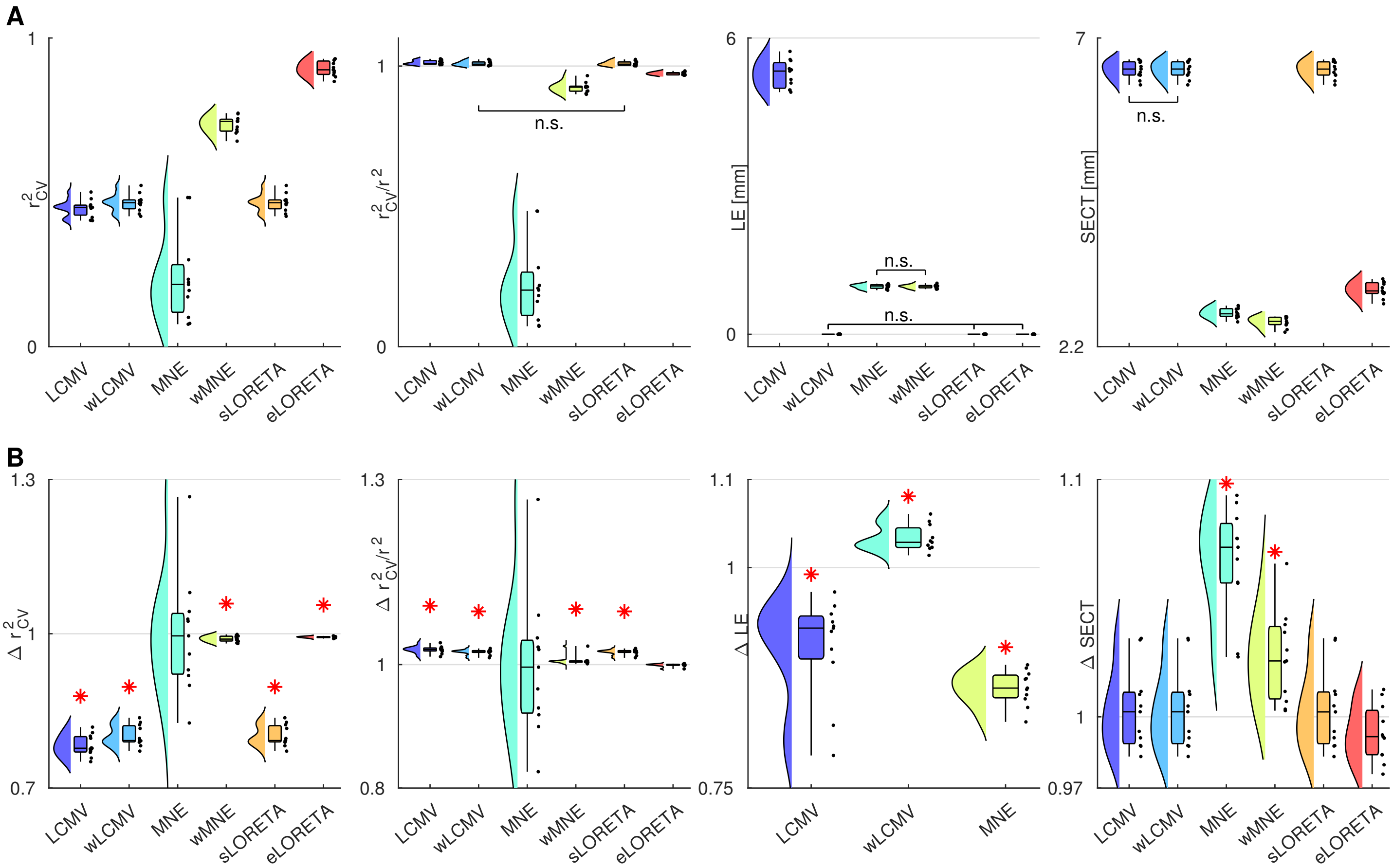


Figure S5: **Comparison between orientation constraints.** (A) Cross-validated variance explained, robustness to cross-validation, LE, and SECT for each algorithm in the eigendecomposition case. For all figures, group-wise analyses demonstrated a significant effect of algorithms. Pairwise analyses identified significant differences for all pairs except those marked as non-significant (n.s.), following false discovery rate correction. (B) Each of the measures in A, shown as a ratio of eigendecomposition to anatomical orientation constraints. Algorithms for which there was a significant pairwise difference between orientation constraints following false discovery rate correction are marked by an asterisk. Values for whiskers, boxes, etc. are as described in Figure 3.

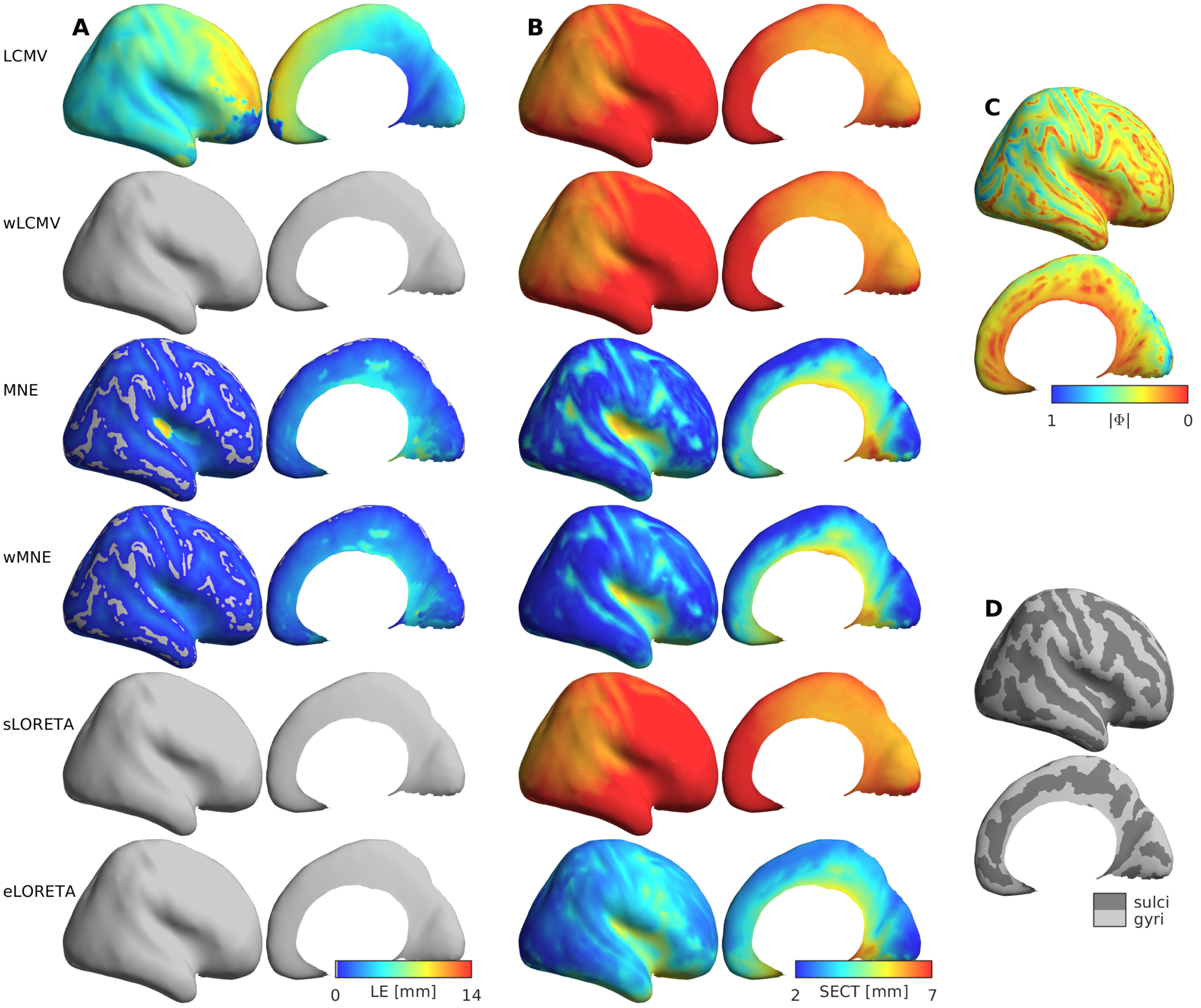


Figure S6: **Spatial distributions of resolution metrics with eigendecomposition orientation constraints.** All plots match those of Figure 4, except eigendecomposition was used for orientation constraints in construction of the leadfield and spatial filter.
